# Physics-based generative model of curvature sensing peptides; distinguishing sensors from binders

**DOI:** 10.1101/2022.09.01.506157

**Authors:** Niek van Hilten, Jeroen Methorst, Nino Verwei, Herre Jelger Risselada

## Abstract

Proteins can specifically bind to curved membranes through curvature-induced hydrophobic lipid packing defects. The chemical diversity among such curvature ‘sensors’ challenges our understanding of how they differ from general membrane ‘binders’, that bind without curvature selectivity. Here, we combine an evolutionary algorithm with coarse-grained molecular dynamics simulations (Evo-MD) to resolve the peptide sequences that optimally recognize the curvature of lipid membranes. We subsequently demonstrate how a synergy between Evo-MD and a neural network (NN) can enhance the identification and discovery of curvature sensing peptides and proteins. To this aim, we benchmark a physics-trained NN model against experimental data and show that we can correctly identify known ‘sensors’ and ‘binders’. We illustrate that sensing and binding are in fact phenomena that lie on the same thermodynamic continuum, with only subtle but explainable differences in membrane binding free energy, consistent with the serendipitous discovery of sensors.

**Teaser:** AI-based design helps explain curvature-selective membrane binding behavior.

## 1 Introduction

The recognition of curved regions of lipid bilayer membranes by proteins plays a key role in many biological processes, such as vesicular transport, fusion, and fission [1, 2]. This preferred binding to curved membranes is called curvature sensing and is driven by the outer leaflet of the curved bilayer membrane being stretched, which causes defects in the packing of the polar lipid head groups. Apolar amino acids of proteins can complement the now exposed hydrophobic tails within these lipid packing defects, negating their energetic penalty and resulting in a thermodynamic driving force (Fig. 1A).

**Fig. 1.**
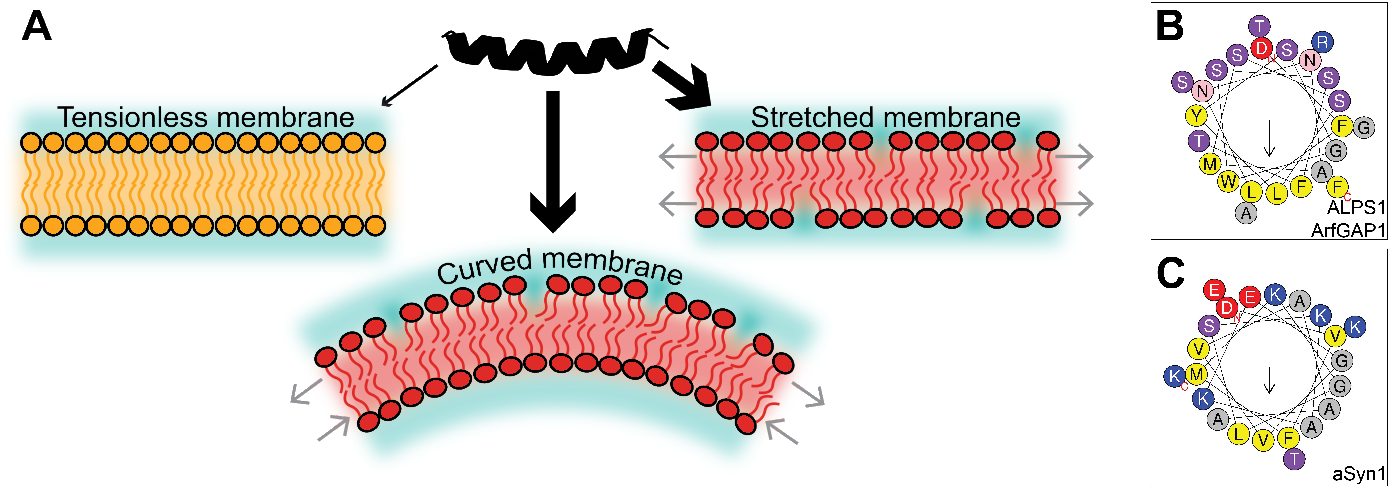
Graphical introduction to curvature sensing peptides. **A)** A peptide (in black) can have an enhanced affinity (thicker arrows) toward a curved or stretched membrane (in red) as opposed to a flat tensionless membrane (in orange), due to hydrophobic lipid packing defects between the lipid head groups. **B-C)** Helical wheel representations [20] of ALPS1 from ArfGAP1 [8] and α-synuclein [9]. Yellow: hydrophobic. Gray: small. Pink/purple: polar. Blue: positively charged. Red: negatively charged.

Besides fundamental biological importance, curvature selectivity has been proposed as a potential avenue for the development of broad-spectrum antiviral peptides that leverage the difference in curvature between the membranes of small enveloped viruses and the essentially flat host cell membrane [3–6]. However, the extremely serendipitous discovery and resulting rarity of curvature selective peptides obstructs the utilization of state-of-the-art data-science driven generative models, like the recent work by IBM on the discovery of antimicrobial peptides [7]. Consequently, an efficient computational strategy for accelerating the discovery of curvature sensing peptides is still lacking.

Many natural curvature sensing proteins feature an amphipathic helix (AH). AHs have a polar face that interacts with the solvent and the lipid head groups and an apolar face that interacts with the hydrophobic lipid tails. Beyond this shared structural amphipathicity, the chemical composition of AHs is highly diverse. For example, the contrasting compositions of the amphipathic lipid packing sensing (ALPS) motif of the ArfGAP1 protein [8] and the AH of *α*-synuclein [9] (Fig. 1B-C) suggest that curvature sensing results from a delicate balance between the amino acid content on the apolar and polar sides of the helices [10, 11]. Moreover, and important to note, some AHs (like *α*-synuclein) have a positive net charge, providing additional selectivity for anionic liposomes specifically [12]. Taken together, the structural diversity among curvature sensors complicates reliable prediction of a given peptide’s sensing ability simply from sequence-based physicochemical descriptors, like mean hydrophobicity 〈*H*〉, hydrophobic moment *μ_H_* [13], and net charge *z*.

Molecular dynamics (MD) simulations are a valuable asset in expanding our understanding of curvature sensing, since they can access the necessary molecular resolution that many experimental methods lack [14–16]. To reduce system size and, consequently, reduce the computational cost, curved membranes are often represented as stretched flat membranes in MD simulations [17, 18] (Fig. 1A), such that the lipid packing defects on the surface are similar and the consequent relative binding free energies correlate [19]. This approximation is valid since biologically relevant bilayer curvatures are negligible on the length scale of peptides (≈5 nm).

Recently, we developed a highly efficient method that can quantify the relative free energy (ΔΔ*F*) of curvature sensing by surface peptides [19]. By redefining ΔΔ*F* as a mechanical property, we realized that differential binding can be reinterpreted as the reduction of work required to stretch a membrane leaflet when the peptide is bound to it. In other words: a peptide that senses leaflet tension/curvature will also tend to induce leaflet tension/curvature and, therefore, those properties are two sides of the same coin. In line with this notion, many AHs have been shown to bind to flat membranes and then actively generate positive membrane curvature (e.g. the AHs of Epsin [21] and the N-BAR domain [22]).

A still unresolved key question is what physical characteristics differentiate AHs that specifically bind to curved membranes (‘sensors’) from AHs that bind without curvature specificity (‘binders’). To date, this question has mostly been addressed by making strategically chosen point mutations in specific example cases [9, 11, 23–25], but a fundamental thermodynamic understanding on how to distinguish ‘sensors’ from ‘binders’ among multiple chemically diverse classes of AHs is – to the best of our knowledge – still lacking. Here, we combine an evolutionary algorithm with coarse-grained molecular dynamics simulations (Evo-MD) to design α-helical peptides that optimally recognize the curvature of lipid membranes, completely from scratch. Our main goal is to demonstrate how the unique synergy between Evo-MD and neural networks can enable the identification of the two major curvature sensing protein families (ALPS and α-synuclein) as well as their known mutants without utilizing any of the available experimental input data, i.e. our approach is purely physics-based. Furthermore, we will illustrate that sensing and binding are in fact phenomena that lie on the same thermodynamic continuum, with only subtle but explainable differences in membrane binding free energy, consistent with the serendipitous discovery of curvature sensors.

## 2 Results

### 2.1 Designing the optimal curvature sensor

Evolutionary molecular dynamics (Evo-MD) is a physics-based inverse design method that embeds molecular dynamics (MD) simulations in a genetic algorithm (GA) framework [26]. GAs are inspired by Darwinian evolution and can serve as a powerful tool for optimization problems in large discrete search spaces, like the 20^L^ possible peptide sequences (for 20 natural amino acids and peptide length *L*). Starting from a, in our case random, initial subset (‘population’), a GA iteratively (1) evaluates the desired property (‘fitness’) of the candidate solutions in the population, (2) selects the best candidates as the ‘parents’ for the next generation, and then (3) performs genetic operations, like cross-over recombination and random point mutations to (4) generate the next population (Fig. 2A). While evolution proceeds, the population’s average fitness will increase until it converges to an optimum. To date, GAs have mainly been applied to peptide optimization problems that involve protein-peptide interactions and use fitness functions based on physicochemical descriptors or information from databases [20, 27–29]. In contrast, the fitness calculation in Evo-MD is based on ensemble averaging from (coarse-grained) MD simulation trajectories and is therefore completely data-independent. In this physics-based approach, experimental data contributes to solving the optimization problem via the parametrization of the force-field that is used in the simulations, the Martini model [30] in this study. Therefore, the main advantage is that Evo-MD will generate curvature sensing peptides without requiring any knowledge of existing curvature sensing peptides, of which too few examples exist to properly train a data-informed model. Additionally, as opposed to data-trained models, physics-based inverse design does not tend to generate molecules that are (too) similar to the input data [31]. In contrast, Evo-MD will search for a pre-defined thermodynamic optimum of sensing and generate physically optimal sequences that actually may differ from the biological optimum due to additional evolutionary constraints imposed by nature’s complexity (e.g. solubility, protein-protein interactions, trafficking).

**Fig. 2.**
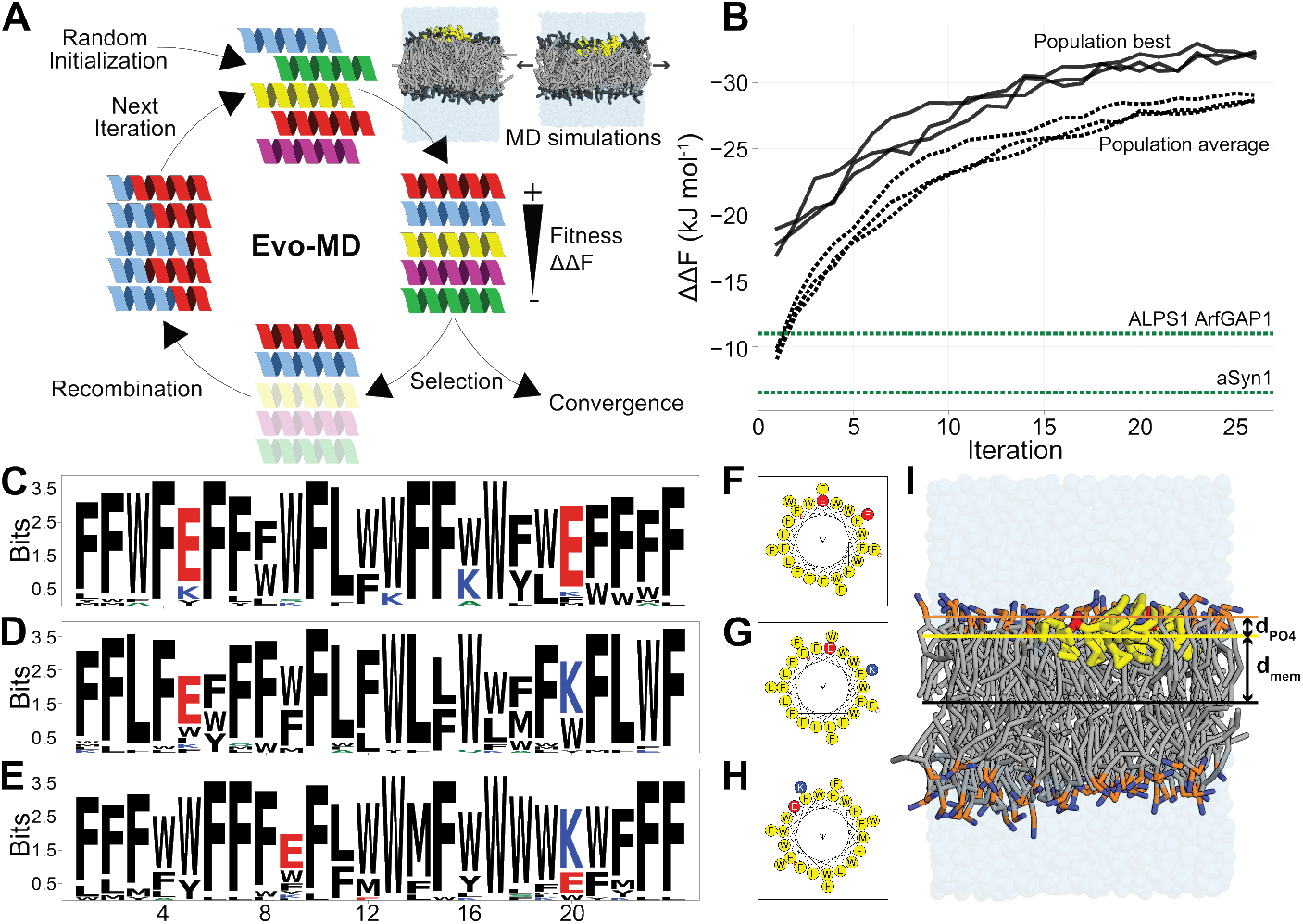
Evo-MD-optimized curvature sensors. **A)** Schematic representation of the Evo-MD process. Figure adapted from previous work [19]. **B)** Three independent replica Evo-MD runs show convergence within 25 iterations, as evident by the population best (solid lines) and population average (dashed lines). For comparison, the ΔΔ*F* values for two known curvature sensing motifs (ALPS1 of ArfGAP1 [8] and *α*-synuclein [9]) are shown in green. We inverted the *y*-axis to emphasize that we maximize the magnitude of ΔΔ*F* whilst retaining the physically relevant minus sign. **C-E)** Consensus sequence logos [40] for the best 36 sequences of the final population. Black: hydrophobic. Blue: positively charged. Red: negatively charged. **F-H)** The respective helical wheel representations [20] of the consensus sequences shown in C-E. Yellow: hydrophobic. Blue: positively charged. Red: negatively charged. **I)** Simulation snapshot of a consensus peptide (Fig. 2C, F) bound to a tensionless POPC membrane. Hydrophobic residues (F, W, L) are shown in yellow; E is shown in red. Phosphate (PO_4_) and choline (NC_3_) beads are shown in orange and blue, respectively. d_mem_ is the *z*-component of the center-of-mass distance between the membrane and the peptide. *d*_PO4_ is the *z*-component of the center-of-mass distance between the PO_4_ groups and the peptide.

The direction of simulated evolution by GAs is governed by the definition of the fitness function (the desired property). For the optimization of curvature sensing peptides, we aim to maximize the magnitude of the curvature sensing free energy ΔΔ*F* that we can efficiently quantify using aforementioned mechanical free-energy method [19]. Our fitness function is the product of the ΔΔ*F* value and a scaling factor c that equals 1 when located on the membrane surface and goes to 0 for transmembrane or soluble configurations (SM section 1). We emphasize that ΔΔ*F* characterizes the relative affinity for lipid packing defects or equivalently positive leaflet curvature, analogous to the curvature-dependent binding constants (free energy of partitioning) measured in experimental model liposome assays [11, 23–25, 32–36] as well as analogous to the measured differences in the concentration of peripheral membrane proteins due to curvature-driven sorting in micropipette aspiration assays [37, 38].

Since membrane-surface peptides are in most cases *α*-helical and the Martini force-field is unable to model protein folding events, we assume and fix helical secondary structure when generating the starting conformations for our peptides. To this end, and to reduce the search space, we excluded 10 amino acids with low *α*-helical propensities [39] (P, G, D, R, C, T, N, H, V, and I), whilst ensuring that every chemical subtype is represented in our final subset (comprising A, L, M, K, Q, E, W, S, Y, and F). We chose a fixed peptide length of 24 residues, which is in the typical range for curvature sensing peptides. Consequently, our search space contains 10^24^ peptide sequences.

In the three independent Evo-MD runs we performed, we observed convergence within 25 iterations with the best candidates having a ΔΔ*F* of around −32 kJ mol^−1^ (Fig. 2B). The consensus sequence logos [41] of the final generations show a strong enrichment of bulky hydrophobic residues, mainly F and W (Fig. 2C-E and supplementary movies). We can understand this result by returning to our earlier statement that “a peptide that senses tension/curvature will also tend to induce tension/curvature”. Such induction of tension and concomitant membrane curvature in a membrane leaflet occurs via shallow insertions within the hydrophobic interior of the lipid membrane, i.e. the region directly below the head groups. Indeed, we observe that the optimal sequences – the point of maximum leaflet tension generation with respect to the helix’ central axis – is characterized by an insertion of 1.69±0.05 nm from the bilayer center in a tensionless membrane (*d*_mem_ in Fig. 2I), or alternatively 0.24±0.05 nm below the average position of the phosphate groups (*d*_PO4_ in Fig. 2I). This is in quantitative agreement with predictions from membrane elastic theory, which suggest an optimal insertion of about 1.7 nm from the membrane center plane [42]. Furthermore, the bulkier the peptide is, and thus the larger its excluded volume and effective helical radius, the more pronounced the induced leaflet tension will be.

Besides the abundant hydrophobic residues, we observed that the solutions of all three Evo-MD runs feature two charged residues (E or K, see Fig. 2C-E). This numerical conservation of two charged amino acids suggests that this is the bare minimum of polar content that is necessary to maintain a surface orientation for such hydrophobic peptides (i.e. scaling factor *c* → 1). The sign of the charge appears to be irrelevant for the zwitterionic POPC membranes we used here. Also, the exact position of these residues seems arbitrary, as long as the two charged residues end up on the same side of the helix in the folded conformation (Fig. 2F-H).

The fact that all three randomly initiated Evo-MD runs produced peptide sequences with identical physical characteristics within the same number of iterations strongly suggests that this is indeed the global and not a local optimum. To probe the effect of only using the 10 most helix-prone amino acids, we performed an additional Evo-MD run with all 20 natural residues included. This, again, yielded peptides with the same physical characteristics, but showing slower convergence (40 iterations) and higher diversity due to the vastly increased search space (Fig. S3).

What this simulated evolution shows is that the GA has successfully selected a key aspect in curvature sensing, insertion of hydrophobic residues [23, 36], which is then maximally amplified and exploited until the fitness converges. To such extent even, that the optimal ‘sensor’ is so hydrophobic that it would likely stick to any membrane, regardless of curvature, thus being classified as a ‘binder’ instead. What is immediately clear is that our optimized peptides strongly differ from the naturally evolved optima (e.g. the ALPS motif and *α*-synuclein), both in terms of ΔΔ*F* (Fig. 2B) and in their chemical compositions (e.g. compare Fig. 1B-C with Fig. 2F-H). Thus, our physics-based inverse design indicates that the distinction between curvature sensors and membrane binders can be considered as a continuum with a soft, subtle threshold at a relative binding free energy that is much lower than the theoretical optimum.

Biologically, the differences between the simulated optimum and naturally evolved peptides can be explained by considering the many boundary conditions imposed by the complex environment of *in vivo* systems. One of the most obvious and fundamental requirements is that proteins should be soluble in physiological buffer. The extremely hydrophobic GA-generated optima clearly fail this criterion and will readily aggregate and precipitate out of the solution. Our finding that curvature sensing is a hydrophobically driven process indicates that natural curvature sensing helical motifs have likely evolved from a trade-off between maximizing hydrophobic insertion (with one face of the helix), whilst remaining generally water-soluble (the other face of the helix), giving rise to their amphipathic character. Also, since curvature sensing implies tension generation, peptides with a high |ΔΔ*F*| could harm the integrity or shape of the membranes they adhere to. To demonstrate this, we performed additional simulations that show that the hydrophobic Evo-MD optimum is sufficiently tension-inducing to generate positive curvature in a flat POPC-membrane (Fig. S4). To circumvent such disruptive effects, an evolutionary pressure to limit this potency must exist.

### 2.2 A neural network model to predict curvature sensing

As a valuable byproduct of the iterative optimization process by Evo-MD, we obtained a large database of ≈54,000 unique sequences (all 24 residues long) and their respective sensing free energies (ΔΔ*F*) as calculated by MD simulations. With this wealth of data, we set out to train a convolutional neural network (CNN) that is able to predict curvature sensing ability from peptide sequence information only.

To enable the model to handle peptides shorter than 24 amino acids as well, we split the sequences in the original data set at a random position, such that the resulting two fragments were at least 7 residues long. Next, since ΔΔ*F* depends linearly on length (Fig. S5), we interpolated the ΔΔ*F* values for the split sequences, hereby tripling the data set to ≈138,000 sequences (after discarding duplicate fragments). We refrained from extrapolating to sequences longer than 24 residues, since this would require additional assumptions on amino acid composition and – potentially – involve more complex tertiary structures that are inaccurately modeled by the Martini force-field. A detailed description of the final training data is included in SM section 6.

As described previously in the context of activity prediction of helical antimicrobial peptides [43], we used one-hot encoded and zero-padded representations for the input sequences. These are then fed to two consecutive convolutional layers with max pooling, followed by a fully connected layer and a single output neuron to translate the connection weights into a float value: the predicted ΔΔ*F* (Fig. 3A, see SM section 7 for details on the optimization of the architecture and hyperparameters).

**Fig. 3.**
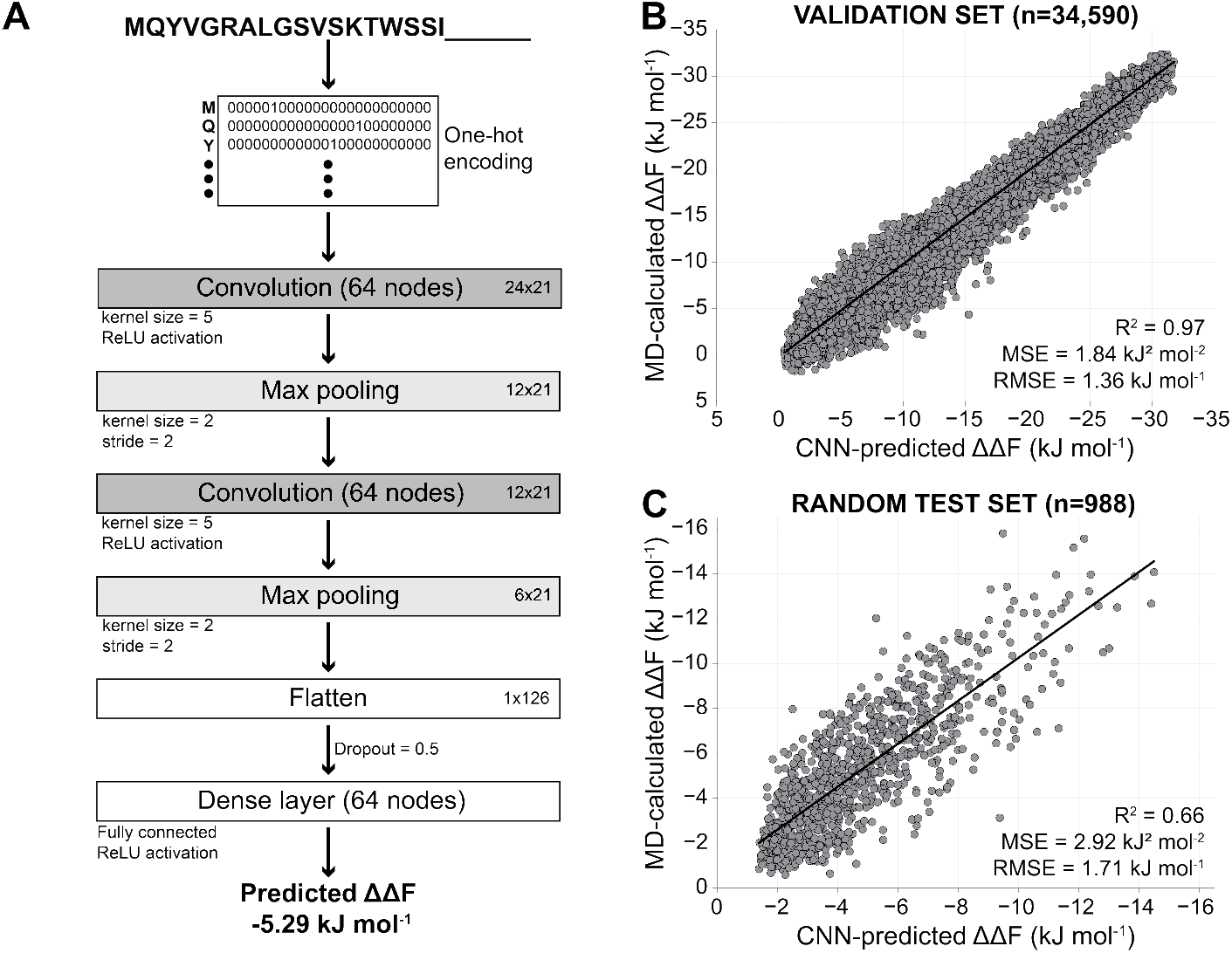
CNN model for curvature sensing prediction. **A)** Architecture of the CNN model. **B-C)** Correlation between the CNN-predicted and MD-calculated ΔΔ*F* values for the validation set after the final training (B) and for a random test set (C).

During the CNN training, minimization of the mean squared error (MSE) converged after 18 epochs (Table S3) to a MSE of 1.84 kJ^2^ mol^−2^ for the validation set (25% random sample from the full data). For this validation set, we achieved excellent correlation (R^2^ = 0.97) between the predicted and MD-calculated ΔΔ*F*’s (Fig. 3B). However, because sequences from the late iterations of the same Evo-MD run can be highly similar, the validation and training sets are arguably not fully independent. Therefore, to ultimately test our model, we predicted ΔΔ*F* for 988 randomly generated sequences (between 7 and 24 residues long) that were not part of the training data and obtained a MSE of 2.92 kJ^2^ mol^−2^ and a *R*^2^-value of 0.66 when comparing the predicted values to ΔΔ*F* calculated by MD simulations (Fig. 3C).

The trained neural network and all data sets are accessible at github.com/nvanhilten/CNN curvature sensing. Please note that the model should only be used for sequences between 7 and 24 amino acids long, and that it assumes α-helical folding (as we did in the training data). Based on the performance of our model on the randomly generated test set, the root-mean-square error (RMSE) of its predictions is 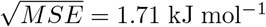, which is comparable to the typical errors obtained when calculating ΔΔ*F* by MD simulation (e.g. compare the error bars in Fig. 4B).

**Fig. 4.**
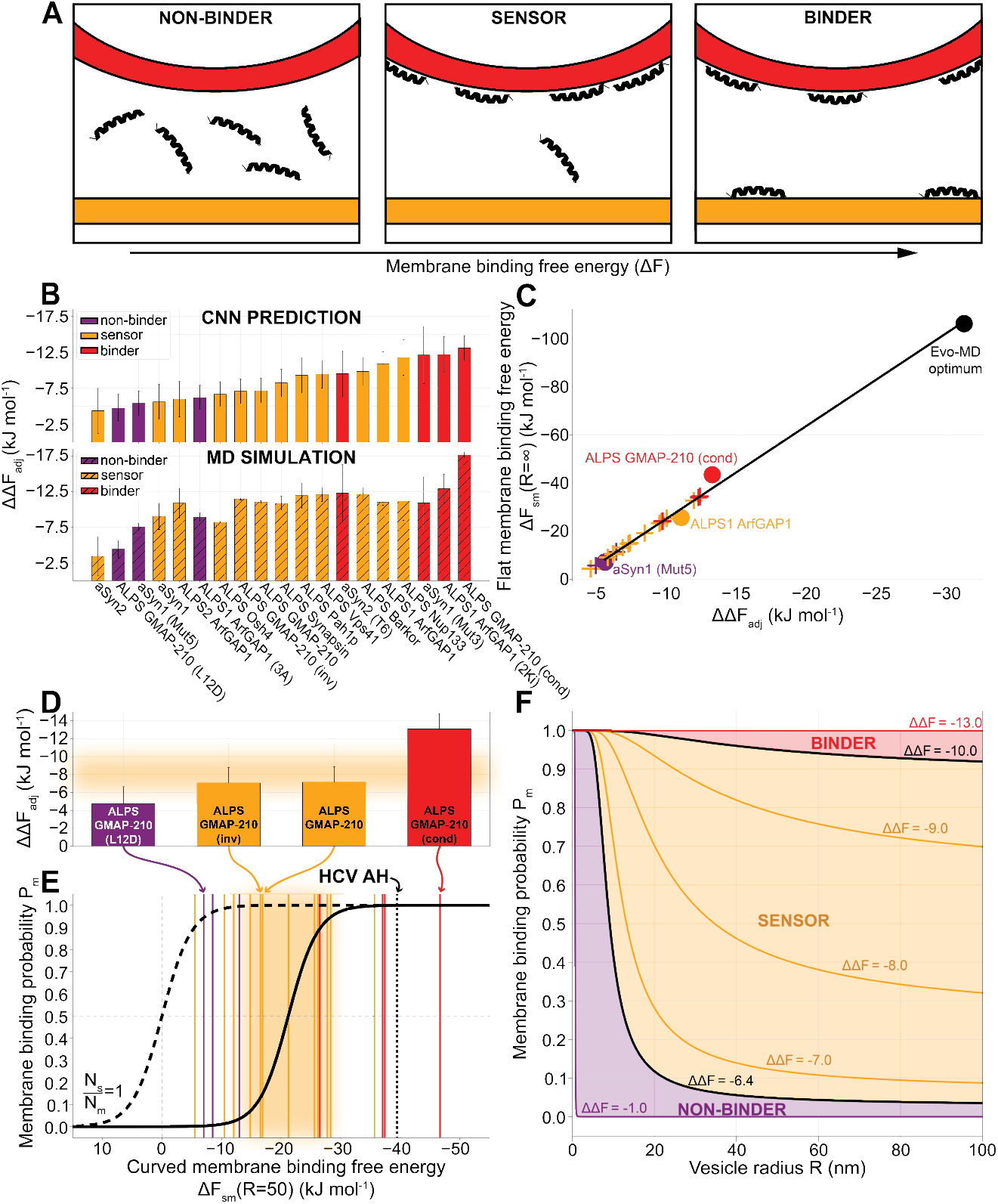
Distinguishing sensors from binders. **A)** “Non-binders” do not bind to membranes; “sensors” only bind to curved membranes; “binders” bind to any membrane, independent of curvature. **B)** ΔΔ*F*_adj_ for 19 benchmark peptides (Table S4) predicted with the CNN model (top) and calculated from MD simulations (bottom). **C)** Linear correlation between CNN-predicted ΔΔ*F*_adj_ and the flat membrane binding free energy Δ*F*_sm_(*R* = ∞). Circles indicate peptides for which Δ*F*_sm_(*R* = ∞) was calculated by thermodynamic integration (SM section 10). The Evo-MD optimum (black) is the sequence in Fig. 2D. For the remaining peptides (crosses), Δ*F*_sm_ was derived from the linear fit Δ*F*_sm_ = 3.83 · ΔΔ*F*_adj_ + 12.27. **D)** Highlighted ΔΔ*F*_adj_ values for ALPS GMAP-210 variants. **E)** The membrane binding probability *P*_m_ as a function of the membrane binding free energy Δ*F*_sm_ (SM section 10 and eq. 2), at a vesicle radius *R* = 50. In D-E, the orange area indicates the likely regime (0.05 ≤ *P*_m_ ≤ 0.95) where peptides are empirically classified as sensors. **F)** Membrane binding probability *P*_m_ as a function of the vesicle radius *R* for a range of sensing free energies ΔΔ*F* (SM section 10).

### 2.3 Distinguishing sensing from binding

Now, with the MD quantification and CNN tools in hand, we can return to the key question posed in the introduction, namely: which, if any, characteristics can help us distinguish curvature sensors from binders, and what can relative binding free energies, like ΔΔ*F*, teach us in this regard? To address this question, we composed a benchmark set of natural curvature sensing peptides (Table S4), also including mutated variants that were empirically categorized as ‘non-binders’ (i.e. no affinity for any membrane) or ‘binders’ (i.e. binding to membranes without curvature specificity). We should acknowledge the expert help of Prof. Bruno Antonny and Dr. Romain Gautier in composing this list. In the following section we will show that these benchmark peptides can be reliably categorized into separate regimes (see Fig. 4A) within a thermodynamic continuum: that is, of a continuous scale in terms of membrane binding free energies (Δ*F*) but features a sharp switch-like transition in terms of membrane partitioning behavior.

To fairly compare the sequences, we propose two correction factors to obtain an adjusted relative binding free energy ΔΔ*F*_adj_. First, we linearly extrapolate the ΔΔ*F* values of shorter peptides to their corresponding free energies if they were 24 residues long (ΔΔ*F*_L=24_, see SM section 5). Second, we realized that many of the peptides are cationic to improve interaction with (curved) anionic membranes that are abundant in nature. Since our MD simulations were performed with neutral POPC membranes, we hypothesized that the relative binding free energies would in these cases be underestimated and thus require a correction term *c*_z_*z* to account for this:

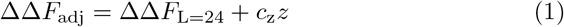

To determine the magnitude of *c*_z_ (the relative free energy contribution per unit charge *z*), we performed additional MD simulations with anionic membranes (75% POPC, 25% POPG) and indeed found elevated relative binding free energies (|ΔΔ*F*_L=24_|), especially for the cationic peptides (Table S5). From the average difference between the ΔΔ*F*’s calculated on the different membranes, we obtained *c*_z_ = −0.93 ± 0.89 kJ mol^−1^ per unit charge.

We calculated ΔΔ*F*_adj_ for the benchmark peptides with the CNN model and MD simulations (Fig. 4B). When ranking the peptides accordingly, we find that we can roughly reproduce the empirical qualification of ‘non-binders’ at the lower end and ‘binders’ at the higher end of the ranking, and ‘sensors’ in the middle. Interestingly, we find that the CNN-predicted ranking is in better agreement with the experimental trends than the results from MD simulations. We speculate that this is likely due to a smoothening effect, i.e. predictions of the MD simulations for individual peptide sequences are fully independent whereas the CNN introduces effective correlations between sequences. Because the CNN is trained on the ensemble averaged values from many thousands of independent MD trajectories for different peptide sequences, we argue that disturbances (‘noise’) in chemical space (point mutations) and in the molecular dynamics itself (limited sampling) are therefore smoothened out to such extent that the experimental trends are more robustly reproduced. Hence, we use the CNN-predicted ΔΔ*F*_adj_ values for the remainder of this paper.

### 2.4 Thermodynamic model of the sensing→binding transition

Along the lines of our current definitions, every peptide undergoing hydrophobically driven membrane binding is able to sense positive membrane curvature due to the increase in surface hydrophobicity upon bending. In other words, positive curvature enhances hydrophobically driven membrane binding. Empirically, however, a peptide is only classified as a curvature sensor if it *only* binds to membranes characterized by a high positive curvature. In this section, we will argue that the empirical classification of sensors versus binders can be intuitively understood from the population statistics of a two-state partition function.

Herein, we define the following two states; (1) state *m*: the peptide is bound to the membrane, (2) state *s*: the peptide is in solution. We define the partitioning free energy difference between the two states as Δ*F*_sm_: the free energy of membrane binding minus the free energy of solvation with respect to the peptide in the gas phase. Using thermodynamic integration, we calculated Δ*F*_sm_ for a non-binder, a sensor, a binder, and the extremely hydrophobic Evo-MD optimum (Fig. 2D, G) binding to a flat tensionless membrane (*R* = ∞, see SM section 10) and found that it linearly relates to ΔΔ*F*_adj_ (Fig. 4C). The reason for this linear correlation is that both membrane binding (Δ*F*_sm_) and curvature sensing (ΔΔ*F*_adj_) are driven by hydrophobic interactions.

Since we are interested in peptides that bind to curved membranes (10 ≤ *R* ≤ 100), we generalize Δ*F*_sm_ to a vesicle radius-dependent form Δ*F*_sm_(*R*) (full derivation in SM section 10) and take a typical radius of *R* = 50 nm for the following calculations.

At thermal equilibrium, the relative probability for a peptide to bind to a membrane *P*_m_ is Boltzmann distributed. Consequently, *P*_m_ is given by:

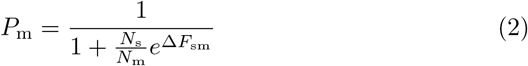

Here, *N*_s_ represents the number of realizations within the accessible solvent and *N*_m_ represents the number of membrane binding realizations. The mathematical form of eq. 2 resembles that of a so-called Fermi-Dirac function. If 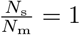, this function features a sharp but continuous transition at Δ*F*_sm_ = 0 (dashed curve in Fig. 4E), i.e. the point where membrane binding and peptide solubility are precisely in balance (*P*_m_ = 0.5). However, the number of realizations in solution is expected to be much larger than the number of realizations associated with membrane binding, 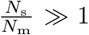, at typical lipid concentrations. Consequently, the transition point shifts to the right, i.e. peptide-membrane binding is favored over peptide solvation at the transition point *P*_m_ = 0.5. The steep nature of this ‘switch function’ strikingly explains *why* empirically classified sensing behavior for one peptide switches to binding behavior for another, possibly highly similar, peptide based on only subtle differences in (relative) membrane binding free energy, like we observed in this work (Fig. 4B).

To finalize our model, we estimate the prefactor 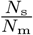. The accessible membrane area in one liter of solution is 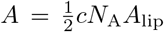, with c being the lipid concentration, *N*_A_ the Avogadro constant, and *A*_lip_ the area per lipid. The characteristic surface area of a helical peptide is roughly *A*_p_ = 5 × 1 nm^2^ and equivalently its volume in solution is *V*_p_ = 5 × 1 × 1 nm^3^. Taking *c* =1 mM (a typical value in the concentration range used in experiments [11, 23–25, 32–36]) and *A*_lip_ = 0.64 nm^2^ we obtain 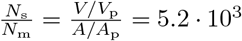.

When we plug this number into eq. 2, we find that the free energy of membrane binding outperforms the free energy of solvation by −21.2 kJ mol^−1^ at the transition point *P*_m_ = 0.5 (Fig. 4E), i.e. the transition point is associated with favorable but relatively weak membrane binding. Because of the sharp transition behavior, ‘sensors’ are envisioned as peptides with a Δ*F*_sm_ value near the transition point (0.05 ≤ *P*_m_ ≤ 0.95, the orange area in Fig. 4E). According to our model, a peptide with *P*_m_ < 0.05 would be classified as a ‘non-binder’, and a peptide with *P*_m_ > 0.95 would be deemed a ‘binder’. When overlaying the values for the 19 benchmark peptides we introduced earlier (vertical lines in Fig. 4E, Table S6) and considering a typical vesicle radius *R* =50 nm, we find that 3/3 non-binders, 8/12 sensors, and 3/4 binders are classified correctly, i.e. in agreement with the original experiments. Such categorization would have been impossible with crude physicochemical descriptors like mean hydrophobicity 〈*H*〉 or hydrophobic moment *μ*_H_ despite them (weakly) correlating with Δ*F*_sm_ (SM section 12).

We defined the sensing regime rather generously (0.05 ≤ *P*_m_ ≤ 0.95) to accommodate for the fact that both the empirical classification and the positioning of the Fermi-Dirac curve are sensitive to the lipid concentration c, which can greatly vary between experiments. Moreover, imperfect helical folding and differing lipid compositions, cell systems, and read-out types can complicate the direct comparison between empirical classifications and our computational models. In terms of the curved membrane binding free energy Δ*F*_sm_(*R* = 50), the lower and upper boundaries of sensing (0.05 ≤ *P*_m_ ≤ 0.95) correspond to −13.9 and −28.5 kJ mol^−1^, respectively (orange area in Fig. 4E). In other words, when the work required to pull a peptide from the membrane of a vesicle with radius *R* = 50 is between those values, it can be classified as a sensor. In terms of the length- and charge-adjusted relative binding free energy (ΔΔ*F*_adj_), i.e. the preference for highly curved membranes (vesicles with radius R=12.5 nm [19]), sensors fall between −6.4 and −10.0 kJ mol^−1^ (orange area in Fig. 4D).

As an example, we highlighted four variants of ALPS GMAP-210 [24, 25] for which the relative differences in experimental sensing/binding behavior were correctly captured (Fig. 4D). This is a striking example because the sequences are similar: the only difference between the original sensor ALPS GMAP-210 and the non-binding variant ALPS GMAP-210 (L12D) is a single point mutation (L → D) that disturbs the peptide’s hydrophobic face. As can be expected, and in line with the experimental findings, the inverse sequence ALPS GMAP-210 (inv) scores the same as the original peptide and is thus categorized as a sensor as well. Finally, the condensed version ALPS GMAP-210 (cond) was correctly identified as a binder. With this highlighted example, we demonstrate that our method and thermodynamic model can (1) pick out features as subtle as single point mutations, (2) resolve the resulting differences in relative free energy, and (3) correctly categorize the consequent sensing behavior.

We also included the antiviral peptide HCV AH, that specifically ruptures vesicles with a high curvature (e.g. small enveloped viruses) [3, 4] (black dotted line in Fig. 4E). To date, HCV AH is the only example of a clinically relevant curvature selective antiviral peptide [5]. When we plug in the previously calculated free energy value [19] for HCV AH, we find that this peptide falls into the ‘binder’ regime. This is consistent with evidence that the vesicle size specificity of this peptide is due to curvature specific pore formation and not to curvature specific binding (i.e. curvature sensing) [44]. After all, subtle binding may not be optimal for pore formation, since the subsequent induction of tension should be sufficient to rupture the membrane.

Following the evaluation of HCV AH in this regard, we argue that the most promising range to find potent curvature specific antiviral agents is therefore near the transition zone between sensing and binding (i.e., *P*_m_ → 1), since these peptides (1) may still benefit from some ‘curvature sensing’ (predominant binding to higher curvatures), (2) pack a larger punch than biological sensors in terms of meeting the tension induction threshold necessary to deform/perforate viral membranes, but are (3) not so potent that they also rupture the host cell membrane. The latter is helped by the fact that the host membrane is likely more resilient to the disruptive actions of peptides than the viral membrane due to membrane stabilizing proteins and active feedback mechanisms. This means that a considerable part of the selectivity of membrane-targeting drugs is likely due to a difference in drug resistance (membrane resilience) rather than actual differential binding.

Finally, we plotted the membrane binding probability Pm as a function of vesicle radius R for different ΔΔ*F* values (Fig. 4F) and find that this is precisely in line with the cartoon in Fig. 4A. Indeed, for non-binders (ΔΔ*F* > −6.4 kJ mol^−1^), membrane binding is rare (low *P*_m_) for biologically relevant vesicle sizes (10 ≤ *R* ≤ 100). Conversely, binders (ΔΔ*F* < −10.0 kJ mol^−1^) have a high membrane binding probability (*P*_m_ → 1) regardless of the vesicle radius. It is exactly the sensor region in between that defines the area where peptides can display curvature selectivity, i.e. a wide range of probabilities for a wide range of vesicle radii.

## 3 Discussion

We have illustrated the utility of a physics-based generative model (Evo-MD) to explore and simultaneously rationalize the mechanisms of how peptides sense membrane curvature. Initially, we set out to optimize curvature sensing by resolving the sequence that maximizes the *relative* affinity for lipid packing defects. Instead, we ended up with the optimal ‘binder’. This finding led to the important realization that curvature sensing and membrane binding are phenomena that lie on the same thermodynamic continuum (Fig. 4E).

Naturally evolved curvature sensors, such as the ALPS motif and α-synuclein, are chemically diverse but turn out to be remarkably similar in terms of partitioning free energies, which explains their functional similarities. In this work, we described the thermodynamic regime that defines the curvature sensing behavior of peptides. Given how narrow this energetic ‘sensing window’ is, it is unsurprising that the discovery and design of curvature sensors has, to date, been rather serendipitous. Having identified this ‘window of opportunity’ in terms of relative binding free energy can facilitate the discovery of previously unidentified curvature sensing peptides since we now know where to look for them.

The existence of the here-resolved thermodynamic sensing regime can also be intuitively understood from an evolutionary biological perspective. Curvature sensing motifs within naturally evolved proteins must fulfil the following two criteria: they should (1) predominantly bind to curved membranes and (2) conserve the structural integrity of the membranes they adhere to. The here-observed linear correlation between sensing and overall membrane binding (Fig. 4C) dictates that these criteria are only met in the weak binding regime. These arguments are all in full agreement with the earlier hypothesis that curvature sensing in nature is a subtle balance between overall membrane binding and specific curvature recognition [2]. Also, in this weak binding regime, the small leaflet strain induced by peptides is able to facilitate a largely inert and thus biologically functional sensing phenotype that does not easily lead to membrane rupture/deformation.

Importantly, we demonstrated a fruitful synergy between a physics-based generative model (Evo-MD) and a convolutional neural network, which not only dramatically accelerated the high-throughput evaluation of peptide sequences, but also improved the accuracy of prediction compared to the original molecular simulations. It is important to stress the key role of Evo-MD, in that it yields sequences over the whole range of ΔΔ*F* by gradually maximizing the relevant chemical property in a well spaced manner, in our example even up to the thermodynamic optimum. Using such data to train a neural network model has the important advantage that it encompasses the full thermodynamic range of possibilities over a vast search space of 20^24^ sequences, whereas a data set of natural peptides (if available in the first place) would be strongly constrained to a certain biologically feasible regime that only comprises peptides with highly similar physicochemical characteristics. We argue that training data generated with Evo-MD can therefore substantially improve both the applicability domain as well as the accuracy of neural network models, despite many of the generated sequences not being necessarily biologically relevant. This principle is equivalent to fitting an unknown function to data points that are well spaced over the whole range of the applicability domain versus data points that are only clustered within a narrow window. Particularly, precise knowledge of the maxima (and minima) of a function – which a physics-based optimization resolves – will benefit the quality of a fit or model, also within the biologically relevant domain of the search space. We postulate that a subsequent restriction of the search space within or near the here-resolved sensing regime can enhance the discovery of curvature sensing motifs in natural proteins, as well as their *de novo* generation.

Finally, we envision an important potential application in the computational design of peptide sensors that recognize membranes with other aberrant characteristics, such as a distinct lipid composition (e.g. bacteria and cancer cells). Since (selective) membrane binding results in the generation of leaflet tension, membrane binding peptides have an inherent membrane destabilizing propensity as well as the ability to lower the energetic cost of the highly curved interface of toroidal pores. This is particularly the case for the hydrophobically driven membrane binding peptides we discussed in this work. The simultaneous encoding of selective membrane binding plus an active drug mode, such as the induction of membrane lysis, is therefore a realistic avenue to explore further. An important advantage of physics-based generative models over existing data-science based generative models herein is their unique ability to systematically explore the drug therapeutic potential of distinct relative binding (ΔΔ*F*) regimes by restricting the generation of peptide sequences within pre-defined boundary values of ΔΔ*F*, for example, via the straightforward introduction of a bias/constraint to the fitness function. This can enable the targeted exploration of different ‘windows of opportunity’ similar to aiming a gun at different targets.

## 4 Materials and Methods

### 4.1 Evo-MD

The Evo-MD code was adapted from previous work [26] to include the preparation, production, and fitness evaluation modules that are specific to the curvature sensing problem. The code is available at github.com/nvanhilten/Evo-MD_curvature_sensing.

Every candidate solution is evaluated by a fitness function that is based on a free-energy calculation from coarse-grained MD simulations, using the Martini 3 force-field [30]. The relative binding free energy (ΔΔ*F*) is calculated from two POPC membrane simulations at constant area: one at tensionless conditions (*σ*(*A*_0_)) and one at stretched conditions (*σ*(*A*\*)) with a relative strain of 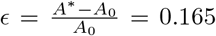, such that the outer leaflet bares lipid packing defects similar to the outside of a highly curved liposome (radius *R* = 12.5 nm). This method is described in detail in previous work [19] and in SM section 1.

To focus the optimization toward surface peptides, we applied a cosine scaling factor c (eq. 3 and SM section 1) that is 1 when the peptide is at the membrane surface (*d*_mem_ = *b*) and 0 when the peptide is in solution (*d*_mem_ ≥ 2b) or fully adopts an inter- or transmembrane configuration (*d*_mem_ ≤ 0). The reference values for *b* are based on the monolayer thickness of a tensionless POPC membrane (*b* = 1.90 nm for *σ*(*A*_0_)) and a stretched POPC membrane (*b* = 1.79 nm for *σ*(*A*\*)) [19] within the Martini 3 force-field. The lowest value for *c* (tensionless or stretched condition) is used in the fitness scaling.

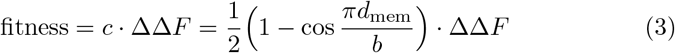

Evo-MD production runs were performed with a population size of 144 peptides, from which a parent pool of the ‘fittest’ 36 sequences was selected. Details on tuning these settings are described in SM section 2. From the parent pool, two parents were selected randomly and combined through one-point crossover recombination: the two parent sequences are split at the same random position *i* and the tail ends (*i* → 24 – *i*) are swapped. Next, for every position in the resulting two ‘children’ peptides, random point mutation is applied with the probability 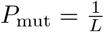, with sequence length *L* = 24, such that every sequence bares – on average – one point mutation.

To retain highly scoring peptides, the best candidate solution of every generation (‘fitness elite’) was copied to the next. Additionally, sequences that were rerun more than 3 times already, were also retained (‘rerun elites’). The fitness values for these elite sequences were updated after every rerun with the average value of the fitness values from all individual iterations. In this way, highly scoring peptides get increasingly better sampling throughout the evolution.

### 4.2 Simulation details

All MD simulations were performed with GROMACS 2019.3 [45] with fixed *x*- and *y*-dimensions (constant area), such that the ‘low tension’ membrane was tensionless (*ϵ* = 0, *A* = *A*_0_) and the ‘high tension’ was at a 16.5% relative strain (*ϵ* = 0.165, *A* = *A*\*) [19]. Berendsen pressure coupling [46] was applied only in the *z*-dimension (1 bar reference pressure with a 4.5 · 10^−5^ bar^−1^ compressibility, *τ*_P_ = 6 ps). A constant temperature of 310 K was maintained by the velocity rescaling thermostat [47] (*τ*_T_ = 1 ps).

We used the coarse-grained Martini force-field, version 3.0.0 [30], at a 30 fs time step. Van der Waals interactions were calculated with the shifted Verlet cut-off scheme [48] and reaction-field electrostatics [49] describe the coulomb potentials, both with a 1.1 nm cut-off. The neighbor list was updated every 20 steps.

Within Evo-MD, the system setup was automated as described in previous work [19]. Systems comprise 128 Martini 16:0–18:1 phosphatidylcholine (POPC) lipids and 1800 Martini water beads. Atomistic helical peptide models were generated using PeptideBuilder [50] and then coarse-grained by martinize2 – VerMoUTH [51]. They were then inserted into the two preequilibrated membrane systems at *d*_mem_ = 1.5 nm from the membrane center plane. For charged peptides, systems were neutralized by adding Na^+^ and Cl^−^ counter ions. Steepest descent minimization with soft-core potentials (0.75 coupling for the Van der Waals interactions) was performed to solve clashes. Within Evo-MD, a total run time of 400 ns was used for each simulation. All non-Evo-MD simulations (Fig. 3C, Fig. 4B, Fig. S5, and Table S5) were run for at least 800 ns.

## Supplementary meterials

- SM section 1, Fig. S1: Fitness function
- SM section 2, Fig. S2: Tuning Evo-MD settings
- SM section 3, Fig. S3: Evo-MD runs including all 20 natural amino acids
- SM section 4, Fig. S4: Curvature generation by Evo-MD optimum
- SM section 5, Fig. S5, Table S1: ΔΔ*F* depends linearly on sequence length
- SM section 6, Fig. S6, Table S2: Training data description
- SM section 7, Table S3: Optimization of neural network architecture and hyperparameters
- SM section 8, Table S4: Benchmark peptide sequences and properties
- SM section 9, Table S5: ΔΔ*F* values for different membrane compositions
- SM section 10, Fig. S7: Calculating Δ*F*_sm_
- SM section 11, Table S6: Classification of benchmark peptides
- SM section 12, Fig. S8: Correlation between Δ*F*_sm_ and Eisenberg parameters
- SM section 13, Movie S1-3: Supplementary movies: evolution of consensus sequences

## Acknowledgments

We thank Prof. Bruno Antonny and Dr. Romain Gautier for insightful discussions and their expert help in selecting relevant curvature sensors, non-binders, and binders to test our theory and methods. We acknowledge the anonymous reviewers for their thorough and considerate remarks, which led to notable improvements of this paper.

## Author contributions

N.v.H. and H.J.R. designed the research. J.M. wrote the Evo-MD code. N.v.H implemented Evo-MD modules specific to the curvature sensing problem. N.v.H. and N.V. developed, trained, and optimized the neural network model. N.v.H. and H.J.R. wrote the manuscript.

## Data avalability

All data needed to evaluate the conclusions in the paper are present in the paper and/or the Supplementary Materials. The Evo-MD and CNN code and datasets are available at zenodo.org/record/7380110.

