## Supplementary information for "Physics-based generative model of curvature sensing peptides; distinguishing sensors from binders"

### Supplementary Materials for *Physics-based generative model of curvature sensing peptides; distinguishing sensors from binders*

Niek van Hilten[1], Jeroen Methorst[1], Nino Verwei[1] and Herre Jelger Risselada[1,2,3*]

[1]Leiden Institute of Chemistry, Leiden University, Einsteinweg 55, Leiden, 2333 CC, The Netherlands.
[2]Department of Physics, Technical University Dortmund, Otto-Hahn-Strasse 4, Dortmund, 44227, Germany.
[3]Institute of Theoretical Physics, Georg-August-University Göttingen, Friedrich-Hund-Platz 1, Göttingen, 37077, Germany.

### Contents

1 **Fitness function** 3

2 **Tuning Evo-MD settings** 5

3 **Evo-MD run including all 20 natural amino acids** 6

4 **Curvature generation by Evo-MD optimum** 7

5 $\Delta\Delta F$ **depends linearly on sequence length** 8

6 **Training data description** 10

7 **Optimization of neural network architecture and hyperparameters** 11

8 **Benchmark peptide sequences and properties** 15

9 $\Delta\Delta F$ **values for different membrane compositions** 16

10 **Calculating** $\Delta F_{\mathrm{sm}}$ 17

11 **Classification of benchmark peptides** 19

12 **Correlation between** $\Delta F_{\mathrm{sm}}$ **and Eisenberg parameters** 20

13 **Supplementary movies: evolution of consensus sequences** 21

### 1 Fitness function

Our fitness function is based on the relative binding free energy $\Delta\Delta F$, that is calculated from the mechanical pathway in the thermodynamic cycle defined in Fig. S1A, as explained in detail in [19]. $\Delta\Delta F$ is the difference in work required to stretch a membrane in presence ($\Delta F_s'$) and absence ($\Delta F_s$) of a peptide bound to its surface. Since the lateral tension $\sigma(A)$ in a membrane is linearly related to the change in membrane area $\frac{A^*-A_0}{A_0}$, the mechanical work of stretching $\Delta F_s = \int_{A_0}^{A^*} \sigma(A)dA$ can be reliably approximated by measuring the surface tension only at two end-states (tensionless $\sigma(A_0)$; and stretched $\sigma(A^*)$) from MD simulations at constant areas $A_0$ and $A^*$. Such that:

$$\begin{aligned} \Delta\Delta F = \Delta F_s' - \Delta F_s &= \int_{A_0}^{A^*} \sigma'(A)dA - \int_{A_0}^{A^*} \sigma(A)dA \\ &= \frac{A^* - A_0}{2}\Big[(\sigma'(A^*) + \sigma'(A_0)) - (\sigma(A^*) + \sigma(A_0))\Big] \end{aligned} \tag{1}$$

The membrane tensions in the peptide-free systems $\sigma(A^*)$ and $\sigma(A_0)$ only have to be measured once to be used in the calculations for any peptide. Consequently, $\Delta\Delta F$ can be calculated from two MD simulations per peptide (to obtain $\sigma'(A^*)$ and $\sigma'(A_0)$), and this is how it was implemented for the fitness calculation in Evo-MD.

We applied a scaling factor $c$ (eq. 2) that is 1 when the peptide is at the membrane surface ($d_{\text{mem}} = b$) and 0 when the peptide is in solution ($d_{\text{mem}} \geq 2b$) or fully adopts an inter- or transmembrane configuration ($d_{\text{mem}} \leq 0$). See Fig. S1B. The reference values for $b$ are based on the monolayer thickness of a tensionless POPC membrane ($b = 1.90$ nm for $\sigma(A_0)$) and a stretched POPC membrane ($b = 1.79$ nm for $\sigma(A^*)$) [19] within the Martini 3 force-field. The lowest value for $c$ (tensionless or stretched condition) is used in the fitness scaling.

$$\text{fitness} = c \cdot \Delta\Delta F = \frac{1}{2}\Big(1 - \cos\frac{\pi d_{\text{mem}}}{b}\Big) \cdot \Delta\Delta F \tag{2}$$

Typically, peptides with a near-optimal $\Delta\Delta F$ (around $-30$ kJ mol$^{-1}$) have values for $c$ of 0.97 and 0.90, for the tensionless and stretched condition, respectively. This corresponds to respective insertion depths ($d_{\text{mem}}$) of 1.69 and 1.42 nm. High values of $c$ mean that the fitness is dominated by $\Delta\Delta F$, as intended. Removing this cosine scaling factor would result in peptide optima that adopt inter- or transmembrane configurations, since such a symmetric embedding in fact results in the maximization of induced tension measured within the constant area ensemble. However, for curvature sensing/induction we are aiming to optimize asymmetric insertion in only the outer leaflet instead [42], thus requiring the scaling factor $c$ to reintroduce this asymmetry in the system.

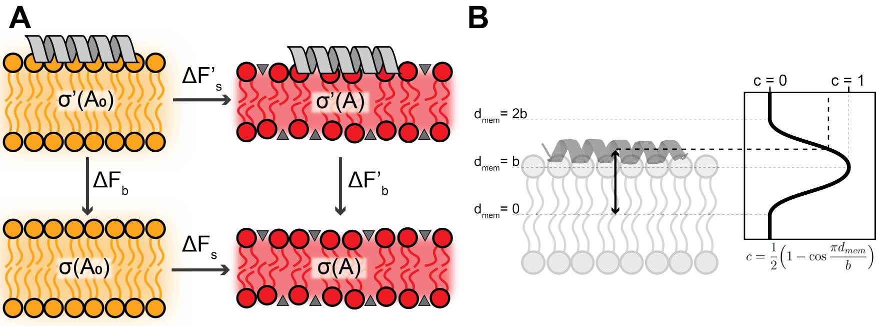


**Fig. S1 Evo-MD fitness function. A)** Thermodynamic cycle that connects the commonly used alchemical pathway (thermodynamic integration of (un)binding, $\Delta F_{\rm b}' - \Delta F_{\rm b}$) to the mechanical pathway (work of stretching, $\Delta F_{\rm s}' - \Delta F_{\rm s}$). Adapted from previous work [19]. **B)** Schematic explanation of the cosine scaling factor $c$, that is related to the insertion depth $d_{\rm mem}$ of the peptide.

### 2 Tuning Evo-MD settings

Evo-MD runs were performed with population sizes ($n$) ranging from 48 to 288. After ranking the population on fitness, the best quarter $p = \frac{n}{4}$ is selected as a parent pool for the next generation. We plotted the convergence (Fig. S2) and decided to perform the production runs with a population size of 144 (and, consequently, 36 parents), since the GA converged to the highest value for these settings (Fig. S2).

Ideally, one would extensively test the effect of changing the GA-related settings (e.g. number of parents, number of elites, mutation rate) on the convergence. However, such optimization is in the case of Evo-MD severely challenged by the computational expense of the fitness calculation by MD simulation (even when using coarse-graining). Taken together, the two MD simulations that are carried out for every peptide require roughly 24 CPU hours to complete on the Intel Cascade Lake Platinum 9242 (CLX-AP) CPUs of the HLRN Lise supercomputer nodes we used. Consequently, every iteration of the GA takes $24 \times 144 = 3,456$ CPU hours. We need at least 20 iterations to reach convergence (Fig. S2), which equates to $3,456 \times 20 = 69,120$ CPU hours for a single GA run. In terms of wall hours, we can run all the simulations of a population in parallel and on 8 cores each ($24/8 = 3$ wall hours per iteration), which means that a converged GA run (20 iterations) requires about 60 wall hours (2.5 days).

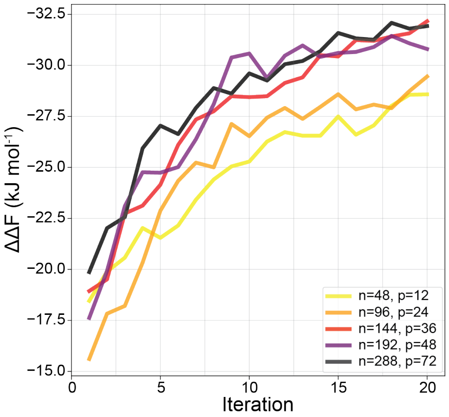


**Fig. S2 Tuning Evo-MD settings.** Convergence of the population's best candidate for different population sizes $n$ and parent pools $p = \frac{n}{4}$

### 3 Evo-MD run including all 20 natural amino acids

In our production runs, we only used the 10 most helix-prone amino acids with all chemical groups represented (small: A; hydrophobic: L, M; polar: S, Q; anionic: E; cationic: K; aromatic: W, Y, and F). To probe the effect of this assumption, we performed an additional Evo-MD run with all 20 natural amino acids (AAs) included.

As can be expected with the vastly increased search space ($10^{24} \rightarrow 20^{24}$), we observed a slower convergence and required the algorithm to run for 40 iterations (Fig. S3A). To test its robustness, we introduced a double mutation rate after 20 iterations, i.e. $P_{\text{mut}} = \frac{2}{L}$ per residue, with $L = 24$. This caused a slight drop in the population's average $|\Delta\Delta F|$ after which it recovered to a similar value of $\approx -25$ kJ mol$^{-1}$. This indicates that the evolution has approached the global optimum, although it is not quite there yet: the best sequences in this 20-AA run score about 2 kJ mol$^{-1}$ lower than the optima from the 10-AA run. However, the general physical characteristics are already fully reproduced, evident from the consensus sequence of the best 36 peptides in the final generation (Fig. S3B) being highly similar to the consensus sequences we found for the 10-AA runs (Fig. 2C-E in main text). As described in the main text, we see a high conservation of big hydrophobic residues (F and W) and a few charged residues that co-locate at the polar side of the amphipathic helix (Fig. S3C). From the reduced height of the characters in the sequence logo, we conclude that the sequences in the 20-AA run are more diverse, due to the larger choice of amino acids and the doubled mutation rate. Generally though, this optimum confirms the conclusions drawn from the original 10-AA runs: the physically optimal curvature sensor is very hydrophobic in nature and would empirically be classified as a binder instead, showing that those two classes are part of the same thermodynamic continuum.

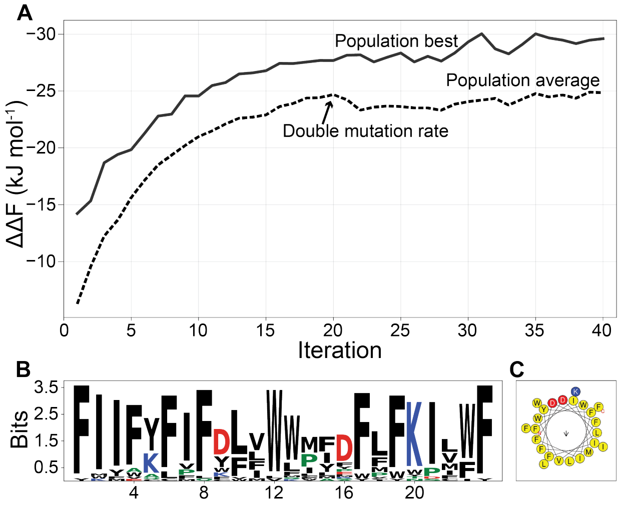


**Fig. S3 Evo-MD run with all 20 natural AAs. A)** The population best (solid line) and population average (dashed line) for a Evo-MD run with all 20 amino acids included. **B)** Consensus sequence logo [40] for the best 36 sequences of the final population. **C)** Helical wheel representation [20] for the consensus sequence in B.

### 4 Curvature generation by Evo-MD optimum

To probe the curvature generative properties of the Evo-MD optimized peptides, we performed a 4 $\mu$s coarse-grained MD simulation of a POPC membrane (441 lipids per leaflet) with and without the hydrophobic consensus peptide (Fig. 2E in main text) bound to the surface. All MD settings were the same as described in section 4.2 of the main text. The local mean curvature in the upper leaflet (to which the peptide was bound) was measured and plotted from the MD trajectory with the membrane-curvature toolkit (`github.com/MDAnalysis/membrane-curvature`) in MDAnalysis [52] (Fig. S4) and showed an induced mean curvature of 0.3 Å$^{-1}$. We should note that the magnitude of the generated curvature is somewhat arbitrary, since it depends on the box area (with periodic boundary conditions). The fact that we find curvature generation for a theoretically optimal curvature sensor shows that those phenomena are indeed two sides of the same coin.

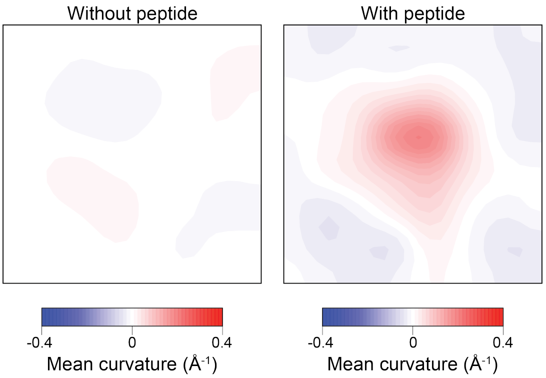


**Fig. S4 Curvature generation by the Evo-MD optimum.** Contour plot of the mean curvature of the upper leaflet of an initially flat 17 × 17 nm$^2$ POPC membrane without a peptide (left) and with a peptide – the Evo-MD optimum `FFFWWFFFEFLWWMFWWWWKWFFF` (Fig. 2E in main text) – bound to its surface. By asymmetrically inducing tension in the upper leaflet (and not in the lower leaflet), peptide inclusion leads to a 0.3 Å$^{-1}$ mean curvature.

### 5 $\Delta\Delta F$ depends linearly on sequence length

Within Evo-MD, we chose a sequence length $L$ of 24 residues, which is a typical length for curvature sensing motifs in natural proteins. To investigate the influence of sequence length on the relative free energy $\Delta\Delta F$, we performed additional calculations for poly-phenylalanine ($F_L$), poly-leucine ($L_L$), poly-alanine ($A_L$), and amphipathic peptides comprising FFQQ repeats (skipping a Q after every 11 residues, ≈3 helical turns, to maximize the hydrophobic moment). We opted to do this analysis for uniform sequences to facilitate the interpretation of the results: any changes in $\Delta\Delta F$ can only be attributed to the change in length, since everything else is the same. F, L, and A were chosen since they are prevalent in $\alpha$-helical motifs and have different hydrophobicities.

The results show that the magnitude of $\Delta\Delta F$ increases linearly with increasing sequence length $L$ (Fig. S5). This finding can be understood from the notion that the relative binding free energy is directly related to the excluded volume of the peptide in the hydrophobic interior of the lipid membrane. Similar to the enrichment of bulky amino acids in the global optima we found by Evo-MD, increasing the sequence length will also increase the total volume of the peptide and, in turn, the magnitude of $\Delta\Delta F$.

We performed a linear fit ($\Delta\Delta F_{\text{fit}}(L) = a_{\text{fit}} L + b_{\text{fit}}$, with $a_{\text{fit}} = -1.03$ and $b_{\text{fit}} = 3.28$, see Fig. S5) on the combined data of poly-F, -L, -A, and -FFQQ. Now, we can use this as a calibration to approximate the relative free energy ($\Delta\Delta F$) of fragments with length $L$ by scaling the calculated $\Delta\Delta F_{\text{L=24}}$ of the full-length original sequence:

$$\Delta\Delta F = \frac{\Delta\Delta F_{\text{fit}}(L)}{\Delta\Delta F_{\text{fit}}(L = 24)} \Delta\Delta F_{\text{L=24}} = \frac{a_{\text{fit}} L + b_{\text{fit}}}{a_{\text{fit}} \cdot 24 + b_{\text{fit}}} \Delta\Delta F_{\text{L=24}} \tag{3}$$

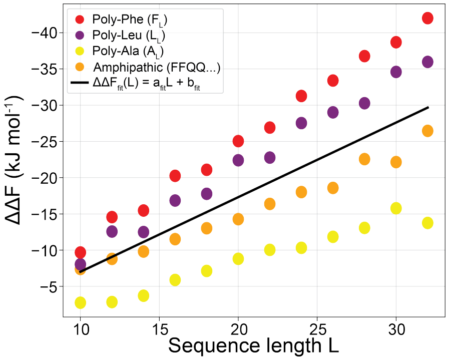


**Fig. S5** **$\Delta\Delta F$ and length relate linearly.** Linear relation between sequence length $L$ and relative free energy $\Delta\Delta F$ for different model sequences.

The physical interpretation of $\Delta\Delta F_{\text{L=24}}$ is the sensing free energy of the peptide *if it were 24 amino acids long*, assuming the same amino acid composition. We validated this length-extrapolation by using the CNN model (see section 2.2 in the main text) to predict $\Delta\Delta F$ for three example ALPS-derived peptides of increasing lengths $(\texttt{DFLNNAMS})_1$, $(\texttt{DFLNNAMS})_2$, and $(\texttt{DFLNNAMS})_3$, and extrapolating the resulting values to $\Delta\Delta F_{\text{L=24}}$ using eq. 3 (Table S1).

**Table S1**: **Example calculation of length-adjusted free energy.** $\Delta\Delta F_{\mathrm{L}=24}$ is the free energy value of a peptide, extrapolated to a sequence length of 24.

| **Sequence** | $\Delta\Delta F$ **(kJ mol$^{-1}$)** | $\Delta\Delta F_{\mathrm{L}=24}$ **(kJ mol$^{-1}$)** |
|---|---|---|
| $(\mathtt{DFLNNAMS})_1$ | -3.674 | -15.881 |
| $(\mathtt{DFLNNAMS})_2$ | -8.341 | -13.548 |
| $(\mathtt{DFLNNAMS})_3$ | -12.928 | -12.928 |
| | **Average** | -14.119 |
| | **SD** | 1.271 |

If both the CNN prediction and the length-extrapolation were perfect, the $\Delta\Delta F_{\mathrm{L}=24}$ values in the right column of Table S1 would be identical. In reality, they have a standard deviation (SD) of 1.271. Since this SD is lower than the expected error of the CNN model (RMSE=1.71 kJ mol$^{-1}$, see Fig. 3C in main text), we conclude from this example that the length extrapolation is valid.

### 6 Training data description

The data set for training our neural network comprises all sequences and respective $\Delta\Delta F$ values we obtained through the Evo-MD tuning (Fig. S2) and production runs (Fig. 2B in main text). A potential issue with training a neural network only on evolutionary optimized data is that such dataset is inherently skewed towards highly scoring sequences. To circumvent this, we performed additional 'steered Evo-MD' runs in which the aim is to minimize the squared distance from a set target value, similar to a harmonic constraint in MD. I.e. the steered Evo-MD fitness (which should be maximized) is given by eq. 4. In total, we performed 20 iterations of such steered Evo-MD (with the same settings as for the main production runs) with $\Delta\Delta F_{\text{target}}$ set to -5, -10, and -15 kJ mol$^{-1}$.

$$\text{fitness} = -(\Delta\Delta F - \Delta\Delta F_{\text{target}})^2 \tag{4}$$

To enable the model to handle all 20 natural amino acids, we also added the results for the 20-AA Evo-MD run (Fig. S3A), as well as 2,500 randomly generated sequences. Finally, sequences from miscellaneous test runs and follow-up projects were included in the data set as well.

**Table S2**: **Composition of the data set.**

| Source | sequences | AAs | Comments |
|---|---|---|---|
| Evo-MD optimization | 12,480 | 10 | Tuning population size (Fig. S2) |
| Evo-MD production runs | 11,232 | 10 | 3 runs of 26 iterations (Fig. 2B) |
| Steered Evo-MD | 8,640 | 10 | $\Delta\Delta F_{\text{target}}$ = -5, -10, or -15 kJ mol$^{-1}$ |
| 20-AA Evo-MD | 5,760 | 20 | 1 run of 40 iterations (Fig. S3A) |
| Random sequences | 2,500 | 20 | |
| Miscellaneous | 14,400 | 10 | Test runs and follow-up projects |
| Miscellaneous | 7,200 | 20 | Test runs and follow-up projects |
| **Total** | 62,212 | | |
| **Total (unique)** | 53,940 | | |

Taken together, the data comprise 53,940 unique sequence-$\Delta\Delta F$ combinations (Table S2), with all peptides being 24 residues long. To also handle shorter peptides – and to expand the training data further – we split all sequences at a random position such that the fragments had a minimal length of 7 amino acids ($\approx$ 2 helical turns). Then, we used the interpolation described previously (Fig. S5 and eq. 3) to obtain an approximated $\Delta\Delta F$ for each of these fragments, and added them to the data set. After removing any newly arisen duplicates, we obtained our final set of 138,358 sequences, the $\Delta\Delta F$ distribution of which is plotted in Fig. S6.

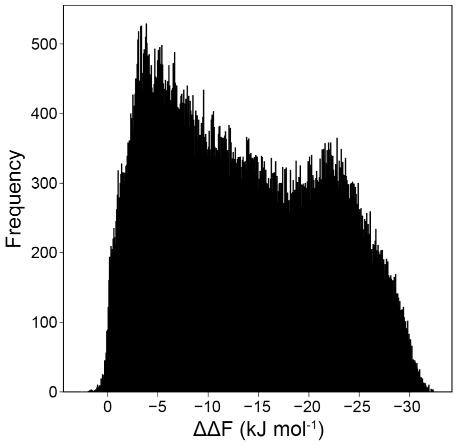


**Fig. S6** **$\Delta\Delta F$ distribution.** $\Delta\Delta F$ distribution of all 138,358 sequences in the data set.

### 7 Optimization of neural network architecture and hyperparameters

25% of the training data was randomly selected as a validation set. The remaining data was used to train the convolutional neural network (CNN), using 5-fold cross validation. Optimization of the CNN was done in two steps, using 988 randomly generated sequences as an independent test set. First, we experimented with different architectures, varying the number of convolutional layers ($N_{\mathrm{c}} \in \{1, 2\}$), the number of nodes per convolutional layer ($N_{\mathrm{n,c}} \in \{32, 64, 128\}$), the number of dense layers ($N_{\mathrm{d}} \in \{1, 2\}$), the number of nodes per dense layer ($N_{\mathrm{n,d}} \in \{32, 64, 128\}$), and the random dropout ($d \in \{0.1, 0.3, 0.5\}$) that is applied when the convoluted and flattened data enters the first dense layer. A batch size ($b$) of 64 and a learning rate ($lr$) of 0.001 were used.

Second, for the best architecture thus far (Fig. 3A in main text, green row in Table S3), we varied the batch size ($b \in \{32, 64, 128\}$) and learning rate ($lr \in \{0.01, 0.001, 0.0001\}$), but found no futher improvement with respect to the original settings ($b = 64$, $lr = 0.001$)

For every tested configuration of the model, we report the average epoch of MSE convergence during cross validation and the final MSE for $\Delta\Delta F$ prediction using the randomized test set (988 sequences, see Fig. 3C in main text) in Table S3. Some models with 32 nodes in the convolutional layer(s) showed no convergence, i.e. the MSE for the validation set did not improve after initialization. We found that dropout had the largest influence on the model's performance, $d = 0.5$ being optimal to prevent over-training.

**Table S3**: **Hyperparameter testing results.** Overview of the tested architectures and hyperparameters, sorted by MSE (in kJ$^2$ mol$^{-2}$). The settings indicated in green were used for the final model.

| $N_c$ | $N_{n,c}$ | $N_d$ | $N_{n,d}$ | **d** | **b** | **lr** | **epoch** | **MSE** |
|---|---|---|---|---|---|---|---|---|
| 2 | 64 | 1 | 64 | 0.5 | 64 | 0.001 | 18 | 2.92 |
| 2 | 32 | 1 | 64 | 0.5 | 64 | 0.001 | 8 | 2.94 |
| 1 | 64 | 1 | 32 | 0.5 | 64 | 0.001 | 62 | 2.96 |
| 1 | 64 | 1 | 128 | 0.5 | 64 | 0.001 | 32 | 3.00 |
| 1 | 32 | 2 | 128 | 0.5 | 64 | 0.001 | 20 | 3.01 |
| 1 | 32 | 2 | 64 | 0.5 | 64 | 0.001 | 43 | 3.05 |
| 1 | 64 | 2 | 128 | 0.5 | 64 | 0.001 | 36 | 3.09 |
| 1 | 32 | 2 | 32 | 0.3 | 64 | 0.001 | 39 | 3.11 |
| 1 | 32 | 1 | 64 | 0.3 | 64 | 0.001 | 42 | 3.13 |
| 1 | 32 | 2 | 32 | 0.5 | 64 | 0.001 | 40 | 3.15 |
| 1 | 64 | 2 | 64 | 0.5 | 64 | 0.001 | 46 | 3.21 |
| 1 | 32 | 1 | 128 | 0.5 | 64 | 0.001 | 33 | 3.22 |
| 2 | 64 | 1 | 64 | 0.5 | 128 | 0.001 | 32 | 3.28 |
| 2 | 64 | 1 | 64 | 0.5 | 32 | 0.01 | 9 | 3.29 |
| 2 | 128 | 2 | 128 | 0.5 | 64 | 0.001 | 19 | 3.30 |
| 1 | 64 | 1 | 64 | 0.5 | 64 | 0.001 | 46 | 3.31 |
| 1 | 32 | 2 | 64 | 0.3 | 64 | 0.001 | 38 | 3.33 |
| 1 | 32 | 1 | 64 | 0.5 | 64 | 0.001 | 38 | 3.34 |
| 1 | 32 | 1 | 32 | 0.3 | 64 | 0.001 | 64 | 3.36 |
| 1 | 64 | 1 | 128 | 0.3 | 64 | 0.001 | 27 | 3.37 |
| 1 | 64 | 1 | 64 | 0.3 | 64 | 0.001 | 42 | 3.37 |
| 2 | 32 | 2 | 64 | 0.3 | 64 | 0.001 | 6 | 3.38 |
| 2 | 32 | 2 | 64 | 0.5 | 64 | 0.001 | Not converged | - |
| 2 | 32 | 1 | 128 | 0.3 | 64 | 0.001 | Not converged | - |
| 2 | 32 | 1 | 128 | 0.5 | 64 | 0.001 | Not converged | - |
| 1 | 128 | 1 | 32 | 0.5 | 64 | 0.001 | 52 | 3.41 |
| 2 | 128 | 2 | 128 | 0.3 | 64 | 0.001 | 19 | 3.44 |
| 1 | 64 | 2 | 64 | 0.3 | 64 | 0.001 | 38 | 3.44 |
| 1 | 64 | 2 | 32 | 0.5 | 64 | 0.001 | 51 | 3.44 |
| 2 | 64 | 1 | 64 | 0.3 | 64 | 0.001 | 25 | 3.45 |
| 2 | 64 | 1 | 64 | 0.5 | 128 | 0.0001 | 108 | 3.45 |
| 2 | 64 | 1 | 128 | 0.3 | 64 | 0.001 | 17 | 3.45 |
| 1 | 64 | 2 | 32 | 0.3 | 64 | 0.001 | 34 | 3.45 |
| 2 | 32 | 1 | 128 | 0.1 | 64 | 0.001 | 22 | 3.46 |
| 1 | 32 | 1 | 32 | 0.5 | 64 | 0.001 | 57 | 3.47 |
| 2 | 64 | 1 | 64 | 0.5 | 64 | 0.01 | 5 | 3.49 |
| 1 | 32 | 1 | 32 | 0.1 | 64 | 0.001 | 40 | 3.49 |
| 2 | 64 | 2 | 64 | 0.5 | 64 | 0.001 | 22 | 3.51 |
| 1 | 128 | 2 | 128 | 0.5 | 64 | 0.001 | 27 | 3.51 |
| 1 | 32 | 2 | 128 | 0.3 | 64 | 0.001 | 33 | 3.52 |
| 2 | 32 | 2 | 128 | 0.1 | 64 | 0.001 | 10 | 3.55 |
| 2 | 64 | 1 | 64 | 0.5 | 32 | 0.001 | 21 | 3.55 |
| 2 | 32 | 1 | 32 | 0.1 | 64 | 0.001 | 26 | 3.56 |
| 1 | 32 | 1 | 128 | 0.3 | 64 | 0.001 | 31 | 3.59 |
| 2 | 128 | 2 | 64 | 0.3 | 64 | 0.001 | 20 | 3.60 |
| 2 | 32 | 1 | 64 | 0.3 | 64 | 0.001 | 17 | 3.60 |
| 1 | 64 | 1 | 32 | 0.3 | 64 | 0.001 | 45 | 3.60 |
| 1 | 128 | 2 | 64 | 0.5 | 64 | 0.001 | 38 | 3.60 |
| 2 | 64 | 1 | 32 | 0.5 | 64 | 0.001 | 21 | 3.61 |
| 2 | 32 | 2 | 32 | 0.1 | 64 | 0.001 | 25 | 3.63 |
| 2 | 64 | 1 | 32 | 0.3 | 64 | 0.001 | 24 | 3.63 |
| 2 | 32 | 1 | 32 | 0.5 | 64 | 0.001 | 30 | 3.64 |
| 2 | 128 | 1 | 64 | 0.5 | 64 | 0.001 | 19 | 3.65 |

| | | | | | | | | |
|---|---|---|---|---|---|---|---|---|
| 1 | 128 | 1 | 32 | 0.3 | 64 | 0.001 | 44 | 3.65 |
| 1 | 128 | 1 | 64 | 0.5 | 64 | 0.001 | 47 | 3.66 |
| 1 | 64 | 2 | 32 | 0.1 | 64 | 0.001 | 23 | 3.67 |
| 1 | 64 | 2 | 128 | 0.3 | 64 | 0.001 | 25 | 3.70 |
| 2 | 32 | 1 | 64 | 0.1 | 64 | 0.001 | 20 | 3.72 |
| 2 | 128 | 1 | 32 | 0.3 | 64 | 0.001 | 22 | 3.74 |
| 2 | 64 | 1 | 32 | 0.1 | 64 | 0.001 | 19 | 3.75 |
| 1 | 128 | 1 | 128 | 0.5 | 64 | 0.001 | 42 | 3.75 |
| 2 | 128 | 1 | 128 | 0.3 | 64 | 0.001 | 25 | 3.75 |
| 2 | 128 | 2 | 64 | 0.5 | 64 | 0.001 | 30 | 3.76 |
| 2 | 64 | 1 | 64 | 0.5 | 64 | 0.0001 | 87 | 3.76 |
| 1 | 128 | 2 | 32 | 0.5 | 64 | 0.001 | 43 | 3.76 |
| 1 | 32 | 2 | 128 | 0.1 | 64 | 0.001 | 19 | 3.77 |
| 2 | 128 | 1 | 64 | 0.3 | 64 | 0.001 | 17 | 3.78 |
| 2 | 64 | 2 | 32 | 0.1 | 64 | 0.001 | 13 | 3.78 |
| 2 | 64 | 1 | 64 | 0.5 | 32 | 0.0001 | 70 | 3.78 |
| 1 | 32 | 2 | 64 | 0.1 | 64 | 0.001 | 20 | 3.79 |
| 1 | 128 | 1 | 128 | 0.3 | 64 | 0.001 | 30 | 3.79 |
| 2 | 128 | 1 | 128 | 0.5 | 64 | 0.001 | 22 | 3.80 |
| 2 | 64 | 2 | 128 | 0.5 | 64 | 0.001 | 23 | 3.81 |
| 2 | 32 | 1 | 32 | 0.3 | 64 | 0.001 | 32 | 3.82 |
| 2 | 32 | 2 | 64 | 0.1 | 64 | 0.001 | 18 | 3.82 |
| 2 | 128 | 1 | 64 | 0.1 | 64 | 0.001 | 12 | 3.82 |
| 2 | 128 | 1 | 128 | 0.1 | 64 | 0.001 | 14 | 3.83 |
| 1 | 128 | 1 | 64 | 0.3 | 64 | 0.001 | 33 | 3.83 |
| 1 | 64 | 2 | 64 | 0.1 | 64 | 0.001 | 18 | 3.87 |
| 2 | 64 | 2 | 32 | 0.5 | 64 | 0.001 | 30 | 3.89 |
| 1 | 64 | 1 | 128 | 0.1 | 64 | 0.001 | 19 | 3.89 |
| 2 | 32 | 2 | 32 | 0.3 | 64 | 0.001 | 26 | 3.90 |
| 2 | 128 | 2 | 64 | 0.1 | 64 | 0.001 | 14 | 3.91 |
| 2 | 128 | 2 | 32 | 0.5 | 64 | 0.001 | 25 | 3.92 |
| 2 | 128 | 2 | 32 | 0.3 | 64 | 0.001 | 17 | 3.93 |
| 2 | 64 | 2 | 128 | 0.1 | 64 | 0.001 | 18 | 3.93 |
| 1 | 32 | 2 | 32 | 0.1 | 64 | 0.001 | 31 | 3.93 |
| 2 | 128 | 1 | 32 | 0.5 | 64 | 0.001 | 20 | 3.94 |
| 1 | 32 | 1 | 64 | 0.1 | 64 | 0.001 | 30 | 3.94 |
| 1 | 128 | 2 | 128 | 0.3 | 64 | 0.001 | 18 | 3.94 |
| 2 | 64 | 2 | 64 | 0.3 | 64 | 0.001 | 31 | 3.96 |
| 1 | 128 | 2 | 32 | 0.3 | 64 | 0.001 | 34 | 3.97 |
| 1 | 128 | 2 | 64 | 0.1 | 64 | 0.001 | 14 | 4.02 |
| 2 | 64 | 1 | 128 | 0.1 | 64 | 0.001 | 17 | 4.03 |
| 1 | 32 | 1 | 128 | 0.1 | 64 | 0.001 | 18 | 4.03 |
| 2 | 64 | 1 | 128 | 0.5 | 64 | 0.001 | 14 | 4.04 |
| 1 | 128 | 1 | 64 | 0.1 | 64 | 0.001 | 29 | 4.07 |
| 1 | 128 | 1 | 128 | 0.1 | 64 | 0.001 | 15 | 4.09 |
| 1 | 128 | 1 | 32 | 0.1 | 64 | 0.001 | 34 | 4.09 |
| 1 | 128 | 2 | 128 | 0.1 | 64 | 0.001 | 11 | 4.12 |
| 1 | 64 | 1 | 32 | 0.1 | 64 | 0.001 | 39 | 4.13 |
| 2 | 64 | 2 | 32 | 0.3 | 64 | 0.001 | 30 | 4.13 |
| 2 | 64 | 2 | 128 | 0.3 | 64 | 0.001 | 33 | 4.14 |
| 1 | 128 | 2 | 32 | 0.1 | 64 | 0.001 | 25 | 4.19 |
| 2 | 128 | 2 | 32 | 0.1 | 64 | 0.001 | 14 | 4.27 |
| 2 | 64 | 1 | 64 | 0.1 | 64 | 0.001 | 18 | 4.29 |
| 2 | 64 | 1 | 64 | 0.5 | 128 | 0.01 | 8 | 4.33 |
| 2 | 32 | 2 | 32 | 0.5 | 64 | 0.001 | 16 | 4.33 |
| 1 | 64 | 1 | 64 | 0.1 | 64 | 0.001 | 33 | 4.38 |
| 2 | 128 | 2 | 128 | 0.1 | 64 | 0.001 | 12 | 4.45 |

| | | | | | | | | |
|---|---|---|---|---|---|---|---|---|
| 1 | 128 | 2 | 64 | 0.3 | 64 | 0.001 | 25 | 4.47 |
| 2 | 128 | 1 | 32 | 0.1 | 64 | 0.001 | 16 | 4.49 |
| 2 | 32 | 2 | 128 | 0.3 | 64 | 0.001 | Not converged | - |
| 2 | 32 | 2 | 128 | 0.5 | 64 | 0.001 | Not converged | - |
| 1 | 64 | 2 | 128 | 0.1 | 64 | 0.001 | 13 | 5.15 |
| 2 | 64 | 2 | 64 | 0.1 | 64 | 0.001 | 18 | 5.44 |

### 8 Benchmark peptide sequences and properties

**Table S4**: **Benchmark peptides.** Sequences, physicochemical properties, and experimental classification of 19 natural benchmark peptides (or derivatives thereof) from the chemically diverse ALPS- and $\alpha$-synuclein (aSyn) families.

| Peptide | Sequence | Length $L$ | Charge $z$ | $\langle H \rangle$ | $\mu_H$ | Exp. classification |
|---|---|---|---|---|---|---|
| ALPS Synapsin | GGGFFSSLSNAVKQTTAAAAATFS | 24 | +1 | 0.373 | 0.328 | Sensor [53] |
| ALPS2 ArfGAP1 | VSQLASKVQGVGSKGWRDVTTFFS | 24 | +2 | 0.370 | 0.441 | Sensor [32] |
| ALPS Kes1/Osh4 | MSQYASSSSWTSFLKSIASFNGD | 23 | 0 | 0.413 | 0.241 | Sensor [23] |
| ALPS Barkor | SSAGGMISSAAASVTSWFKAYTG | 23 | +1 | 0.439 | 0.344 | Sensor [33] |
| ALPS Vps41 | FNPTTNIGSLLSSAASSFRGTPDK | 24 | +1 | 0.310 | 0.365 | Sensor [34] |
| ALPS Nup133 | LPQGQGMLSGIGRKVSSLFGILS | 23 | +2 | 0.555 | 0.417 | Sensor [54] |
| ALPS Pah1p | MQYVGRALGSVSKTWSSI | 18 | +2 | 0.476 | 0.557 | Sensor [35] |
| ALPS GMAP-210 | MSSWLGGLGSGLGQSLGQVGGSLA | 24 | 0 | 0.536 | 0.437 | Sensor [24] |
| ALPS GMAP-210 (inv) | LSGGVQGLSQGLGSGLGGLWSSML | 24 | 0 | 0.594 | 0.424 | Sensor [25] |
| ALPS GMAP-210 (L12D) | MSSWLGGLGSGDGQSLGQVGGSLA | 24 | -1 | 0.433 | 0.335 | Non-binder [25] |
| ALPS GMAP-210 (cond) | MSSWLGGLLGSLVGSLLTIFKDML | 24 | 0 | 0.823 | 0.466 | Binder [25] |
| ALPS1 ArfGAP1 | DFLNNAMSSLYSGWSSFTTGASRF | 24 | 0 | 0.464 | 0.486 | Sensor [8] |
| ALPS1 ArfGAP1 (3A) | DFLNNAMSSAYSGASSATTGASRF | 24 | 0 | 0.263 | 0.306 | Non-binder [23] |
| ALPS1 ArfGAP1 (2Ki) | DFLNNAMSKLYSGWSSFKTGASRF | 24 | +2 | 0.372 | 0.483 | Binder [23] |
| aSyn1 | DVFMKGLSKAKEGVVAAAEKTK | 22 | +2 | 0.129 | 0.354 | Sensor [9] |
| aSyn1 (Mut5) | RVFMKGLSDADRFVVAFARDTD | 22 | 0 | 0.273 | 0.444 | Non-binder [9] |
| aSyn1 (Mut3) | DVFMKGLSKAKAFVVAFAAKTK | 22 | +4 | 0.364 | 0.323 | Binder [9] |
| aSyn2 | KTKQGVAEAAGKTKEGVLYVGSKT | 24 | +3 | 0.064 | 0.215 | Sensor [11] |
| aSyn2 (T6) | KLKQGVAEAAGKFKEGVLYVGSKL | 24 | +3 | 0.248 | 0.374 | Binder [11] |

### 9 $\Delta\Delta F$ values for different membrane compositions

Many biological membranes are anionic. Hence, many curvature sensing motifs in natural proteins have evolved to be cationic to improve membrane interactions and potentially enable additional composition sensing. This aspect is not captured by our initial MD simulations, since we use neutral POPC membranes in our setup. To correct for this discrepancy, we performed additional (and otherwise identical) $\Delta\Delta F$ calculations on membranes with 25% of negatively charged 16:0–18:1 phosphatidylglycerol (POPG).

Due to additional coulomb interactions, we generally found higher (more negative) free-energy values on the negatively charged membrane, especially for positively charged peptides (Table S5). On average, we observed a change of $c_z = -0.93 \pm 0.89$ kJ mol$^{-1}$ in $\Delta\Delta F$ for every unit of net charge. I.e. raising the net charge of a peptide by one boosts the magnitude of $\Delta\Delta F$ by $\approx 1$ kJ mol$^{-1}$.

**Table S5**: **Sensing free energies for different membrane compositions.** Length-adjusted $\Delta\Delta F_{\mathrm{L}=24}$ values calculated from MD simulations ($n$ = 3) on anionic (75% POPC, 25% POPG) and neutral (100% POPC) membranes, with standard deviations $\sigma$. The charge correction factor $c_z$ is the average value of the difference (diff) between the two $\Delta\Delta F_{\mathrm{L}=24}$'s devided by the respective peptide charge $z$, and thus only exists for non-neutral peptides. The final column gives the charge-adjusted $\Delta\Delta F_{\mathrm{adj}} = \Delta\Delta F_{\mathrm{L}=24} + c_z z$ (eq. 1 in main text). The standard deviation $\sigma_{c_z}$ of the diff/$z$ values for all non-neutral peptides is propagated to the final error as $\sigma_{\mathrm{adj}} = \sqrt{\sigma^2 + (z\sigma_{\mathrm{cz}})^2}$. All free energies are given in kJ mol$^{-1}$.

| | | **75% POPC, 25% POPG** | **100% POPC** | | **100% POPC** |
|---|---|---|---|---|---|
| **Peptide** | $z$ | $\Delta\Delta F_{\mathrm{L}=24}$ | $\Delta\Delta F_{\mathrm{L}=24}$ | diff/$z$ | $\Delta\Delta F_{\mathrm{adj}}$ |
| ALPS Synapsin | +1 | $-10.11 \pm 0.59$ | $-9.94 \pm 0.40$ | −0.17 | $-10.87 \pm 0.98$ |
| ALPS2 ArfGAP1 | +2 | $-12.47 \pm 0.49$ | $-9.01 \pm 1.15$ | −1.73 | $-10.88 \pm 2.12$ |
| ALPS Kes1/Osh4 | 0 | $-7.78 \pm 0.92$ | $-8.22 \pm 0.11$ | - | $-8.22 \pm 0.11$ |
| ALPS Barkor | +1 | $-12.15 \pm 0.31$ | $-11.21 \pm 0.28$ | −0.95 | $-12.14 \pm 0.94$ |
| ALPS Vps41 | +1 | $-12.99 \pm 0.31$ | $-11.30 \pm 0.41$ | −1.82 | $-12.11 \pm 0.98$ |
| ALPS Nup133 | +2 | $-13.15 \pm 0.64$ | $-9.32 \pm 2.07$ | −1.92 | $-11.19 \pm 2.73$ |
| ALPS Pah1p | +2 | $-13.67 \pm 0.22$ | $-10.06 \pm 0.14$ | −1.80 | $-11.93 \pm 1.79$ |
| ALPS GMAP-210 | 0 | $-12.94 \pm 0.33$ | $-11.04 \pm 0.19$ | - | $-11.04 \pm 0.19$ |
| ALPS GMAP-210 (inv) | 0 | $-13.22 \pm 0.24$ | $-11.47 \pm 0.14$ | - | $-11.47 \pm 0.14$ |
| ALPS GMAP-210 (L12D) | -1 | $-6.51 \pm 0.85$ | $-5.36 \pm 0.86$ | 1.15 | $-4.42 \pm 1.24$ |
| ALPS GMAP-210 (cond) | 0 | $-20.04 \pm 0.13$ | $-17.64 \pm 0.42$ | - | $-17.64 \pm 0.42$ |
| ALPS1 ArfGAP1 | 0 | $-12.58 \pm 0.25$ | $-10.99 \pm 0.39$ | - | $-10.99 \pm 0.39$ |
| ALPS1 ArfGAP1 (3A) | 0 | $-10.26 \pm 0.17$ | $-8.92 \pm 0.59$ | - | $-8.92 \pm 0.59$ |
| ALPS1 ArfGAP1 (2Ki) | +2 | $-13.00 \pm 0.41$ | $-11.05 \pm 0.92$ | −0.97 | $-12.91 \pm 2.01$ |
| aSyn1 | +2 | $-8.14 \pm 0.34$ | $-7.18 \pm 0.33$ | −0.48 | $-9.04 \pm 1.82$ |
| aSyn1 (Mut5) | 0 | $-7.58 \pm 0.59$ | $-7.57 \pm 0.55$ | - | $-7.57 \pm 0.55$ |
| aSyn1 (Mut3) | +4 | $-13.63 \pm 0.96$ | $-7.18 \pm 0.63$ | −1.61 | $-10.91 \pm 3.63$ |
| aSyn2 | +3 | $-3.26 \pm 0.57$ | $-0.66 \pm 0.56$ | −0.87 | $-3.46 \pm 2.74$ |
| aSyn2 (T6) | +3 | $-9.53 \pm 1.79$ | $-9.48 \pm 2.98$ | −0.02 | $-12.28 \pm 4.01$ |
| | | | $c_z$ | −0.93 | |
| | | | $\sigma_{c_z}$ | 0.89 | |

#### 10 Calculating $\Delta F_{\text{sm}}$

The partitioning free energy difference $\Delta F_{\text{sm}}$ between the membrane-bound state $m$ and solvation $s$ was determined through thermodynamic integration (TI) [55] (Fig. S7). The system setup of the peptide-membrane system is the same as the tensionless system ($R = \infty$) that was implemented in Evo-MD. Now, however, simulations were run with semiisotropic pressure coupling rather than at constant area. The peptide-solvent system was prepared by removing the lipid molecules from the system, performing a steepest decent minimization, and a 50 ns MD simulation to obtain an equilibrated $NpT$ ensemble. The Van der Waals- and Coulomb-interactions between peptide particles and their surrounding were gradually switched off (coupling parameter $\lambda = 0 \rightarrow \lambda = 1$ with 37 intermediate states), such that the fully decoupled configuration is the peptide in vacuum. For the peptide-membrane system, we used a soft harmonic constraint ($k_{\text{force}} = 50$ kJ mol$^{-1}$ nm$^{-2}$) to retain a membrane bound conformation in high decoupling states, as described previously [19]. We probed the influence of this harmonic constraint by performing additional TI simulations for one of the curvature sensors (ALPS1 ArfGAP1) with a five fold higher force constant ($k_{\text{force}} = 250$ kJ mol$^{-1}$nm$^{-2}$), which yielded the same value for $\Delta F_{\text{sm}}(R = \infty)$: $-26.271 \pm 0.342$ kJ mol$^{-1}$, compared to $-25.670 \pm 0.988$ kJ mol$^{-1}$ for the original setting ($k_{\text{force}} = 50$ kJ mol$^{-1}$nm$^{-2}$). This shows that the contribution of this constraint to the membrane binding free energy is negligible. We also tried to perform additional TI calculations with a five fold weaker force constant ($k_{\text{force}} = 10$ kJ mol$^{-1}$nm$^{-2}$), which destabilized the simulations to such extent – especially in the high decoupling states – that we were unable to calculate a $\Delta F_{\text{m}}$ value for this setting. This demonstrates the importance of introducing a weak constraint to prevent the peptide from "flying across the box". The latter would render the free energy difference between the interacting and non-interacting state dependent on the system's dimensions, whereas $\Delta F_{\text{sm}}$ in our model is a scale-independent intrinsic property which is independent of the system's dimensions.

The Langevin stochastic dynamics (SD) integrator and thermostat were used for the TI runs. Every $\lambda$-state was run for 500 ns, with the first 50 ns being discarded from the analysis for equilibration. Free energies were obtained through numerical integration over the ensemble averaged $\frac{\partial V(\lambda)}{\partial \lambda}$ values, with $V$ being the potential energy:

$$\Delta F_{\text{sm}} = \Delta F_{\text{s}} - \Delta F_{\text{m}} = \left[ \int_0^1 \left\langle \frac{\partial V(\lambda)}{\partial \lambda} \right\rangle_\lambda d\lambda \right]_{\text{s}} - \left[ \int_0^1 \left\langle \frac{\partial V(\lambda)}{\partial \lambda} \right\rangle_\lambda d\lambda \right]_{\text{m}} \quad (5)$$

Since we are interested in peptides that bind to curved membranes ($10 \leq R \leq 100$), we need to rewrite $\Delta F_{\text{sm}}$ as a function of the vesicle radius $R$, which includes an offset given by the difference in binding free energy between a flat tensionless membrane ($R = \infty$) and a curved membrane with radius $R$; i.e. $\Delta\Delta F(R)$:

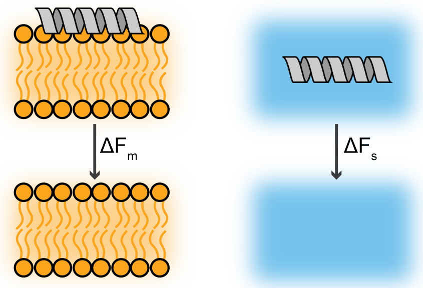


**Fig. S7 Thermodynamic integration.** Schematic representation of thermodynamic integration to obtain the membrane binding free energy $\Delta F_{\mathrm{sm}}(R=\infty)$ between the peptide binding to a tensionless membrane (left) and the solvation energy (right).

$$\Delta F_{\mathrm{sm}}(R) = \Delta F_{\mathrm{sm}}(R=\infty) + \Delta\Delta F(R) \tag{6}$$

Throughout this paper, we calculate $\Delta\Delta F(R)$ for a set relative strain $\epsilon = 0.165$. As we have derived in our previous work [19], $\epsilon$ relates to the vesicle radius $R$, with $d$ being half the monolayer thickness, thus $d \simeq 1$ in for our 4 nm thick POPC bilayer:

$$\epsilon(R) = \frac{d^2}{R^2} + \frac{2d}{R} = \frac{1}{R^2} + \frac{2}{R} \tag{7}$$

This relation yields a radius of $R = 12.5$ nm for our default calculation of $\Delta\Delta F(R)$ with $\epsilon(R) = 0.165$.

Because $\Delta\Delta F(R)$ and $\epsilon(R)$ are linearly related (for relative strains below $\epsilon(R=12.5) = 0.165$, see Fig. 3C in our previous work [19]), we can generalize for all $R$ from $\Delta\Delta F(R=12.5)$:

$$\Delta\Delta F(R) = \frac{\Delta\Delta F(R=12.5)}{\epsilon(R=12.5)}\epsilon(R) = \frac{\Delta\Delta F(R=12.5)}{0.165}\left(\frac{1}{R^2} + \frac{2}{R}\right) \tag{8}$$

When we plug eq. 8 into eq. 6, we obtain:

$$\Delta F_{\mathrm{sm}}(R) = \Delta F_{\mathrm{sm}}(R=\infty) + \frac{\Delta\Delta F(R=12.5)}{0.165}\left(\frac{1}{R^2} + \frac{2}{R}\right) \tag{9}$$

Because the curvature stress vanishes with $1/R^2$, the smaller the vesicles become, the greater the offset $\Delta\Delta F(R)$ will influence the overall membrane binding free energy $\Delta F_{\mathrm{sm}}(R)$ due to the increase of hydrophobic lipid packing defects on the surface. For typical liposome sizes in experiments (e.g. $R = 50$), the contribution of $\Delta\Delta F(R)$ is rather small, since $\frac{1}{50^2} + \frac{2}{50} = 0.04$.

### 11 Classification of benchmark peptides

To validate our method, we compared the predictions of our physics-trained CNN model ($\Delta\Delta F_{\mathrm{adj}}$, and related $P_{\mathrm{m}}$) with experimental classifications from the literature for the 19 ALPS- and aSyn-related example peptides. 3/3 non-binders, 8/12 sensors, and 3/4 binders are correctly classified.

**Table S6**: **Classification of the benchmark peptides.** Benchmark peptides sorted by $\Delta\Delta F_{\mathrm{adj}}$ (predicted by the CNN, Fig. 4B in main text). $\Delta F_{\mathrm{sm}}(R = 50)$ is the binding free energy for vesicles with a 50 nm radius, as calculated by eq. 9 in SM section 11. $P_{\mathrm{m}}$ is calculated from $\Delta F_{\mathrm{sm}}(R)$ using eq. 2 in the main text. Errors are propagated from the obtained MSE of the CNN model on the randomized test set with the error of the charge correction $z\sigma_{\mathrm{c_z}}$, such that $\sigma_{\mathrm{adj}} = \sqrt{MSE + (z\sigma_{\mathrm{c_z}})^2}$. Free energies are given in kJ mol$^{-1}$. Non-binder: $P_{\mathrm{m}} < 0.05$; Sensor: $0.05 \leq P_{\mathrm{m}} \leq 0.95$; Binder: $P_{\mathrm{m}} > 0.95$.

| Peptide | $\Delta\Delta F_{\mathrm{adj}}$ | $\Delta F_{\mathrm{sm}}(R = 50)$ | $P_{\mathrm{m}}$ | Pred. classification | Exp. classification | Correct? |
|---|---|---|---|---|---|---|
| aSyn2 | $-4.36 \pm 3.18$ | $-5.49$ | 0.00 | Non-binder | Sensor [11] | no |
| ALPS GMAP-210 (L12D) | $-4.73 \pm 1.93$ | $-6.99$ | 0.00 | Non-binder | Non-binder [25] | yes |
| aSyn1 (Mut5) | $-5.38 \pm 1.71$ | $-8.51$ | 0.01 | Non-binder | Non-binder [9] | yes |
| aSyn1 | $-5.59 \pm 2.47$ | $-10.50$ | 0.01 | Non-binder | Sensor [9] | no |
| ALPS2 ArfGAP1 | $-5.97 \pm 2.47$ | $-12.04$ | 0.02 | Non-binder | Sensor [32] | no |
| ALPS1 ArfGAP1 (3A) | $-6.20 \pm 1.71$ | $-12.98$ | 0.03 | Non-binder | Non-binder [23] | yes |
| ALPS Osh4 | $-6.65 \pm 1.71$ | $-14.81$ | 0.07 | Sensor | Sensor [23] | yes |
| ALPS GMAP-210 (inv) | $-7.07 \pm 1.71$ | $-16.52$ | 0.13 | Sensor | Sensor [25] | yes |
| ALPS GMAP-210 | $-7.16 \pm 1.71$ | $-16.89$ | 0.15 | Sensor | Sensor [24] | yes |
| ALPS Synapsin | $-8.23 \pm 1.93$ | $-21.25$ | 0.50 | Sensor | Sensor [53] | yes |
| ALPS Pah1p | $-9.30 \pm 2.47$ | $-25.60$ | 0.85 | Sensor | Sensor [35] | yes |
| ALPS Vps41 | $-9.44 \pm 1.93$ | $-26.17$ | 0.88 | Sensor | Sensor [34] | yes |
| aSyn2 (T6) | $-9.52 \pm 3.18$ | $-26.50$ | 0.89 | Sensor | Binder [11] | no |
| ALPS Barkor | $-9.83 \pm 1.93$ | $-27.76$ | 0.93 | Sensor | Sensor [33] | yes |
| ALPS1 ArfGAP1 | $-10.85 \pm 1.71$ | $-28.33$ | 0.95 | Sensor | Sensor [8] | yes |
| ALPS Nup133 | $-11.77 \pm 2.47$ | $-35.66$ | 1.00 | Binder | Sensor [54] | no |
| aSyn1 (Mut3) | $-12.11 \pm 3.93$ | $-37.05$ | 1.00 | Binder | Binder [9] | yes |
| ALPS1 ArfGAP1 (2Ki) | $-12.20 \pm 2.47$ | $-37.41$ | 1.00 | Binder | Binder [23] | yes |
| ALPS GMAP-210 (cond) | $-13.09 \pm 1.71$ | $-46.69$ | 1.00 | Binder | Binder [25] | yes |

### 12 Correlation between $\Delta F_{\text{sm}}$ and Eisenberg parameters

The Eisenberg parameters mean hydrophobicity $\langle H \rangle$ and hydrophobic moment $\mu_{\text{H}}$ [13] are often used to characterize amphipathic helical peptides. For a given sequence, $\langle H \rangle$ and $\mu_{\text{H}}$ can be calculated from a hydrophobicity scale [56] of the individual amino acids. $\langle H \rangle$ is simply the average value of the individual hydrophobicities in the sequence. $\mu_{\text{H}}$ is the mean vector sum of the individual hydrophobicities, such that it is a measure for the segregation of polar and apolar residues in a helical orientation.

We plotted the partitioning free energy between a curved membrane and solvation $\Delta F_{\text{sm}}(R = 50)$ against $\langle H \rangle$ and $\mu_{\text{H}}$ for our set of example peptides and found that they both weakly correlate (Fig. S8). Despite this correlation, reliable ranking and classification like we have done with our free-energy based thermodynamic Fermi-Dirac model (Fig. 4 in main text) would have been impossible with the crude (but more readily calculated) Eisenberg descriptors $\langle H \rangle$ and $\mu_{\text{H}}$.

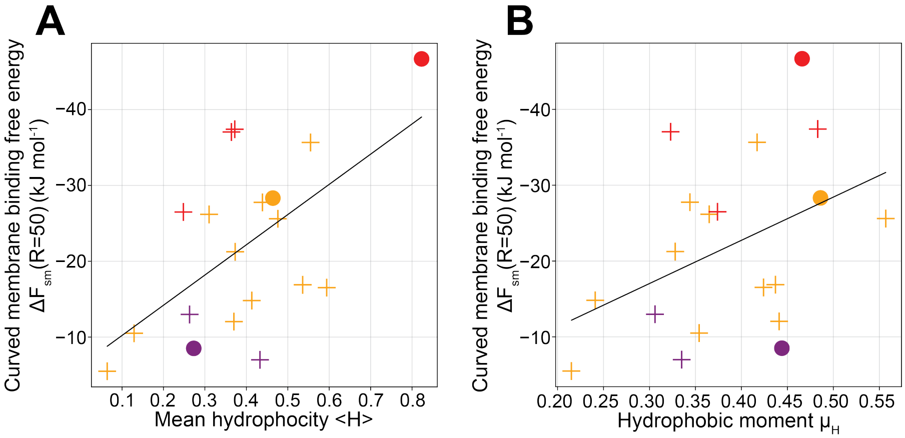


**Fig. S8 Correlation between membrane binding free energy and Eisenberg parameters. A)** Correlation between the curved membrane binding free energy $\Delta F_{\text{sm}}(R = 50)$ and mean hydrophobicity $\langle H \rangle$. **B)** Correlation between $\Delta F_{\text{sm}}(R = 50)$ and hydrophobic moment $\mu_{\text{H}}$.

### 13 Supplementary movies: evolution of consensus sequences

Supplementary movie S1-3 show the evolution of the consensus sequence logos over the course of three independent Evo-MD runs (Fig. 2B in main text). The final frames in movie S1-3 correspond to the consensus sequences in Fig. 2C-E, respectively.
